# Genetics of tibia bone properties of crossbred commercial laying hens in different housing systems

**DOI:** 10.1101/2021.06.21.449243

**Authors:** Martin Johnsson, Helena Wall, Fernando A Lopes Pinto, Robert H. Fleming, Heather A. McCormack, Cristina Benavides-Reyes, Nazaret Dominguez-Gasca, Estefania Sanchez-Rodriguez, Ian C. Dunn, Alejandro B. Rodriguez-Navarro, Andreas Kindmark, Dirk-Jan de Koning

## Abstract

Osteoporosis and bone fractures are a severe problem for the welfare of laying hens, with genetics and environment, such as housing system, each making substantial contributions to bone strength. In this work, we performed genetic analyses of bone strength, bone mineral density and bone composition, as well as body weight, in 860 commercial crossbred laying hens from two different companies, kept in either furnished cages or floor pens. We compared bone traits between housing systems and crossbreds, and performed a genome-wide association study of bone properties and body weight.

As expected, the two housing systems produced a large difference in bone strength, with layers housed in floor pens having stronger bones. These differences were accompanied by differences in bone geometry, mineralisation and chemical composition. Genome-scans either combining or independently analysing the two housing systems revealed no genome-wide significant loci for bone breaking strength. We detected three loci for body weight that were shared between the housing systems on chromosomes 4, 6 and 27 (either genome-wide significant or suggestive when the housing systems were analysed individually) and these coincide with associations for bone length.

In summary, we found substantial differences in bone strength, content and composition between hens kept in floor pens and furnished cages that could be attributed to greater physical activity in pen housing. We found little evidence for large-effect loci for bone strength in commercial crossbred hens, consistent with a highly polygenic architecture for bone strength in the production environment. The lack of consistent genetic associations between housing systems in combination with the differences in bone phenotypes support gene-by-environment interactions with housing system.

## Introduction

Osteoporosis and bone fractures, and more generally poor bone quality, are a severe problem for the welfare of laying hens, with genetics and environment, such as housing system, each making substantial contributions to bone strength. Over their lifetimes, layers experience progressive weakening of the structural bone (Cransberg et al., 2001; Wilson et al., 1992) and increasing risk of fractures. The heritability of tibiotarsal breaking strength, one of the main phenotypes used to measure bone strength, is estimated to be around 0.2-0.5 (Bishop et al., 2000; González-Cerón et al., 2015; Mignon-Grasteau et al., 2016).

Housing has a fundamental and complex influence on the bones of layer hens. On the one hand, housing systems that allow for more exercise promote bone development whereas systems that restrict movement induce bone loss, as bone adapts to loading (Aguado et al., 2015; Fleming et al., 2006, 1994; Jendral et al., 2008; Leyendecker et al., 2005; Newman and Leeson, 1998; Rodriguez-Navarro et al., 2018; Shipov et al., 2010). On the other hand, systems that encourage movement may also increase the fracture risk, for example due to accidental fall from height or collision (Abrahamsson and Tauson, 1993; Fleming et al., 2006; Gregory et al., 1990; Hester et al., 2013). Modern furnished cages allow for more movement and have a more complex environment than the non-furnished cages of old, but there are still environmental differences relevant to bone health between furnished cages and non-cage systems (Rodenburg et al., 2008; Wilkins et al., 2011). In commercial flocks housed in aviaries with different complexity, bone strength is higher in the more complex housing systems where hens move more (Pufall et al., 2021). Housing system also affects the geometry, mineralization and composition of bone, with non-caged birds having thicker and more mineralised cortical bone, and a larger amount of medullary bone, suggesting a greater capacity for bone formation in birds that can exercise more (Fleming et al., 2006; Rodriguez-Navarro et al., 2018; Shipov et al., 2010).

The genetic basis of bone strength in laying hens has previously been mapped in experimental intercrosses and within pedigree lines (Dunn et al., 2007; Raymond et al., 2018), but layer hens on-farm are generally crossbred and kept in different housing systems. This may make the genetic architecture of bone strength on-farm different from conditions previously studied by researchers, especially if there is gene-by-environment interaction. In particular, the genes which are involved in bone turnover in response to mechanical stimuli may differ from those involved in bone development in an environment with reduced mobility and bone loading.

In this work, we performed a genome-wide association study of tibial breaking strength, bone content and composition, as well as body weight in 860 commercial crossbred hens from two different companies, kept in either furnished cages or floor pens. We used a three-point bending test, peripheral quantitative computed tomography (QCT) and thermogravimetric analysis (TGA) to estimate differences in bone strength, bone geometry, mineralization, and chemical composition between the housing systems.

## Materials and Methods

### Crossbred layer hens

Crossbred layer hens of the genotypes Bovans White and Lohmann Selected Leghorn Classic (LSL) were reared at the same commercial rearing farm. Pullets destined for housing in floor pens were reared in an aviary system with full access to all tiers. Pullets destined for furnished cages were fenced in one of the tiers of the aviary to resemble rearing in a conventional rearing cage.

### Management and housing

At 15 weeks of age, the pullets were transferred to the poultry experimental facility at the Swedish Livestock Research Centre Lövsta and subsequently housed either in furnished 8-hen cages or in a one-tier floor housing system. The housing systems and management has been described in Wall et al. 2021. The study was performed with ethical approval from the Uppsala Local Ethics Committee. In brief, each furnished cage provided 600 cm^2^ cage are per hen, 150 cm^2^ nest area, 150 cm litter area (on top of the nest box) and 15 cm perch length per hen (Victorsson Industrier AB, Frillesås, Sweden). Twice a week, litter boxes were replenished with saw-dust and manure belts underneath the cage were run. Each floor pen comprised 13.4 m^2^ and was equipped with Vencomatic® one-tier system (Vencomatic Group, Eersel, The Netherlands). Two thirds of the floor area was a raised slatted area where nests, perches, circular feed hoppers and bell drinkers were located. The remaining floor area was covered with wood shaving. Each pen housed 102 layers. Scrapes under the slatted area removed manure twice a week. A lighting schedule providing 9 hours of light per day on arrival, with a successive increase to 14 hours at 23 weeks was applied in both housing systems.

As part of the same study, we evaluated the effect of organic zinc supplementation in feed. The sampled hens were from both dietary treatments (252 treatment and 257 control in furnished cages; 224 treatment and 235 control in floor pens). As the dietary treatment was not significantly associated with bone strength (average difference of 1.7 N, p = 0.54 in a linear model including housing system and crossbred) we did not include diet in any of the further analyses in this paper. A detailed description of the organic zinc supplementation treatment and analyses of its effect on mortality, integument and bone strength will be published in (Wall et al., n.d.).

### Bone phenotyping

At 100 weeks of age, material for bone phenotyping was collected. An intravenous injection of pentobarbital sodium (100mg/ml) euthanized the layers. Body weight was recorded and a necropsy was conducted to make sure that only hens still in lay were chosen for bone phenotyping. The main phenotype for genome-wide association was tibiotarsal breaking strength (load to failure – we refer to it as “bone strength” for the rest of the paper).

Quantitative computerized tomography (QCT) was performed with the Stratec QCT XCT Research M (Norland; v5.4B) operating at a resolution of 70 µm as previously described (Rubin et al., 2007). Trabecular bone mineral density, which in the female bird reflects bone mineral density of both trabecular and medullary bone, was determined *ex-vivo*, with two metaphyseal QCT scans of the region situated at six percent of bone length from the distal end, and the medullary/trabecular bone was defined by setting an inner threshold to density mode (400 mg/cm^3^). In addition to medullary/trabecular bone data, scans of the metaphyseal area were also used for derivation of data for total bone. Cortical bone parameters were determined *ex-vivo* with a mid-diaphyseal QCT scan of the tibia. After the QCT analyses the tibia were stored at –20°C until biomechanical testing was performed.

The tibiotarsal bones, which had previously been measured by QCT, were subsequently tested for biomechanical strength in a three-point bending test on an electromechanical testing machine (Avalon technologies, Rochester, MN, USA). The specimens were kept frozen until a few hours prior to testing when the bones were completely thawed at room temperature. The specimens were placed with the posterior cortex resting against two end supports placed with a distance of 40 mm between them. The bones were placed in such a way that the load was applied 6 mm distal from the mid part of the tibiotarsal diaphysis with an anterio-posterior direction. The aim was to apply the load at the level where QCT measurements had been performed. An axial load cell (Sensotec inc., Columbus, OH, USA) with the range 0-500 N was used to apply a load of one mm/s to the bone. Values for load and displacement were collected 50 times per second until failure using software provided with the testing machine (Testware II). Based on the collected data load at failure was calculated.

Because these QCT phenotypes are highly correlated (Supplementary Figure 1), we used principal component analysis to reduce the QCT data to three principal components that we used for genome-wide association. The first principal component had high loadings for most of the radiographic phentoypes, while the second had high loadings for bone length, and the third for mostly cortical density (Supplementary Figure 2).

We used thermogravimetric analysis to measure bone mineralization and composition (in cortical and medullary bone, separately), and that mainly consist of water, organic matter (collagen), and mineral (carbonate, calcium, phosphate). Powdered bones were treated at 200, 600, and 800 °C in a RWF 1100 furnace (Carbolite, UK) for one hour and weighed to determine the weight fraction of main bone chemical components. We estimated the percentage water (H2O%), organic matrix (organic%), mineral (mineral%) of the bone, as well as the percentage calcium phosphate (PO4%) and carbonate (CO3%) that are the main mineral part components. We calculated the degree of mineralization (PO4/organic) and the relative content of carbonate in the mineral (CO3/PO4). Because the thermogravimetric phenotypes are less correlated than the tomography phenotypes, we analysed them separately instead of trying to reduce them with principal components (Supplementary Figure 3).

The resulting sample sizes for each set of phenotypes are shown in Supplementary Table 1.

The scanning electron microscopy images in Figure 2 were taken from mid diaphyseal cross-sections of the tibiae. Bones were embedded in EpoThin expoxy resin (Buehler), cut, polished and coated with carbon (Hitachi UHS evaporator). They were imaged with FEI Quanta 400 scanning electron microscope using a backscattering electron detector.

### Genotyping

We genotyped 882 hens at 57,636 single nucleotide variants, using the Illumina Infinium assay. The genotyping was performed by the SNP&SEQ Technology Platform at Uppsala University, Uppsala, Sweden. We excluded 14 individuals with high missingness, as well as 19 individuals that appeared to be recorded as the wrong crossbred based on a principal component plot of the genotypes (Supplementary Figure 4). In order to place the SNP markers on the latest reference genome, we aligned sequences flanking the markers to the chicken reference genome version GRCg6a with BLAT (Kent, 2002).

### Comparisons between housing systems

We compared bone phenotypes and body weight between housing systems using linear models including housing system and crossbred as covariates, and then estimated the contrast between housing systems within each crossbred. Thus, the model was:

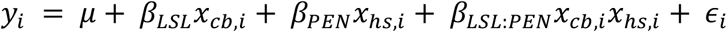

Where *y*_*i*_ is the trait value, *μ* the coefficient for Bovans hens in furnished cages, *β*_*LSL*_ the coefficient for LSL hens, *β*_*PEN*_ the coefficient for floor pens, *β*_*LSL*:*PEN*_ coefficient for the interaction, *x*_*cb,i*_ and *x*_*hs,i*_ indicator variables for crossbreds and housing systems respectively, and *∈*_*i*_ a normally distributed error term. The contrasts of interest were −*β*_*PEN*_, the difference between floor pens and cages within the Bovans crossbreds, and −*β*_*PEN*_ − *β*_*LSL*:*PEN*_, the difference between floor pens and cages within the LSL crossbreds.

We used R statistical environment (R Core Team, 2017), and the *multcomp* package for fitting linear contrasts (Hothorn et al., 2008).

### Genome-wide association studies

We performed genome-wide associations studies using linear mixed models and a genomic relationship matrix, following the approach of (Rönnegård et al., 2016). That is, we first used the *hglm* R package (Rönnegård et al., 2010) to fit a linear mixed model, and use the covariance structure for this model and ordinary least squares to fit the model for each marker efficiently.

We performed genome scans separately for each housing system and jointly, combining the housing systems. Bone phenotype scans included body mass and crossbred, and in the case of joint scans also housing system, as fixed factors. Body weight scans included crossbred, and in the joint scan also housing system, as fixed factors. Genome scans of floor pens included the pen group as a random effect. Joint scans included group as a random effect, combining all furnished cages into one dummy group. We used a conventional genome-wide significance threshold of 5 * 10^−8^, and a suggestive threshold of 10^−4^. Supplementary Dataset 1 contains the summary statistics for all markers.

We used the same linear mixed models to estimate genomic heritability explained by the genomic relationship matrix, and perform a likelihood ratio test against a model without the additive genetic effect as a significance test of the heritability.

### Bivariate genomic models

We used GCTA to estimate genomic heritability and genomic correlations between bone breaking strength in the two different housing systems (Lee et al., 2012), using breed and body weight as fixed effects. The software fits a bivariate linear mixed model using the genomic relationship matrix:

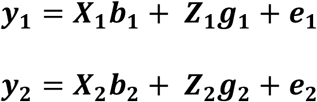

where ***y***_**1**_and ***y***_**2**_ are vectors of trait values; ***b***_**1**_ and ***b***_**2**_ are vectors of coefficients for the fixed effects (breed and body weight); ***g***_**1**_ and ***g***_**2**_ are vectors of additive genetic effects, ***X***_**1**_, ***X***_**2**_, ***Z***_**1**_ and ***Z***_**2**_; ***e***_**1**_ and ***e***_**2**_ are residuals. The variance—covariance matrix uses the genomic relationship matrix derived from genotypes.

### Attempted replication of previously detected bone loci

We attempted to replicate associations from genome-wide association and linkage mapping studies of bone traits from a pedigree line and an experimental intercross (Johnsson et al., 2015; Raymond et al., 2018). The selected candidate regions are listed in Supplementary Table 2. We used genome-wide association summary statistics from markers within 50 kbp of these regions.

### Overlap with previously published loci from chicken QTLdb

We used the GALLO R package (Fonseca et al., 2020) to perform a enrichment test with known quantitative trait loci from the Chicken QTLdb database (Hu et al., 2015) and a hypergeometric test. We mapped the QTL coordinates from the chicken reference genome version Galgal5.0 to GRCg6a with the UCSC LiftOver tool, which resulted in a total of 8427 QTL that could be mapped.

### Availability of data and code

The summary statistics of all genome-wide association studies are included in the paper as Supplementary Dataset 1.

The underlying data have been deposited to Figshare with doi 10.6084/m9.figshare.14405894, containing one file of SNP chip genotypes; one phenotype file of bone traits, body weight and covariates; a file mapping phenotype column names to the trait names used in the article; and one file of marker positions.

The analysis scripts are available at https://github.com/mrtnj/layer_bone_gwas.

## Results

### Differences between housing systems and crossbreds

As expected, bone strength (load to failure) was higher (on average 65 N) in the floor pen system than in the cage system, while body weight was similar. Figure 1 shows body weight and tibial breaking strength in both housing systems and crossbreds, with estimated differences from a linear model. The crossbreds had similar tibial breaking strength, but Bovans were on average 55 g heavier than LSL hens. Figure 2 displays electron microscopy images of tibia from hen housed in a floor pen and a hen housed in a furnished cage showing the distribution of cortical and medullary bone in cross-section. The hen housed in a floor pen had a thicker cortex and a larger amount of medullary bone than the hen from a furnished cage. Also, medullar bone particles are larger and interconnected in the floor pen whereas in furnished cage particles are smaller and isolated. These differences suggest that hens housed in floor pens have a greater capacity to form bone and mineralise the medullar cavity than hens housed in furnished cages.

**Figure 1.**
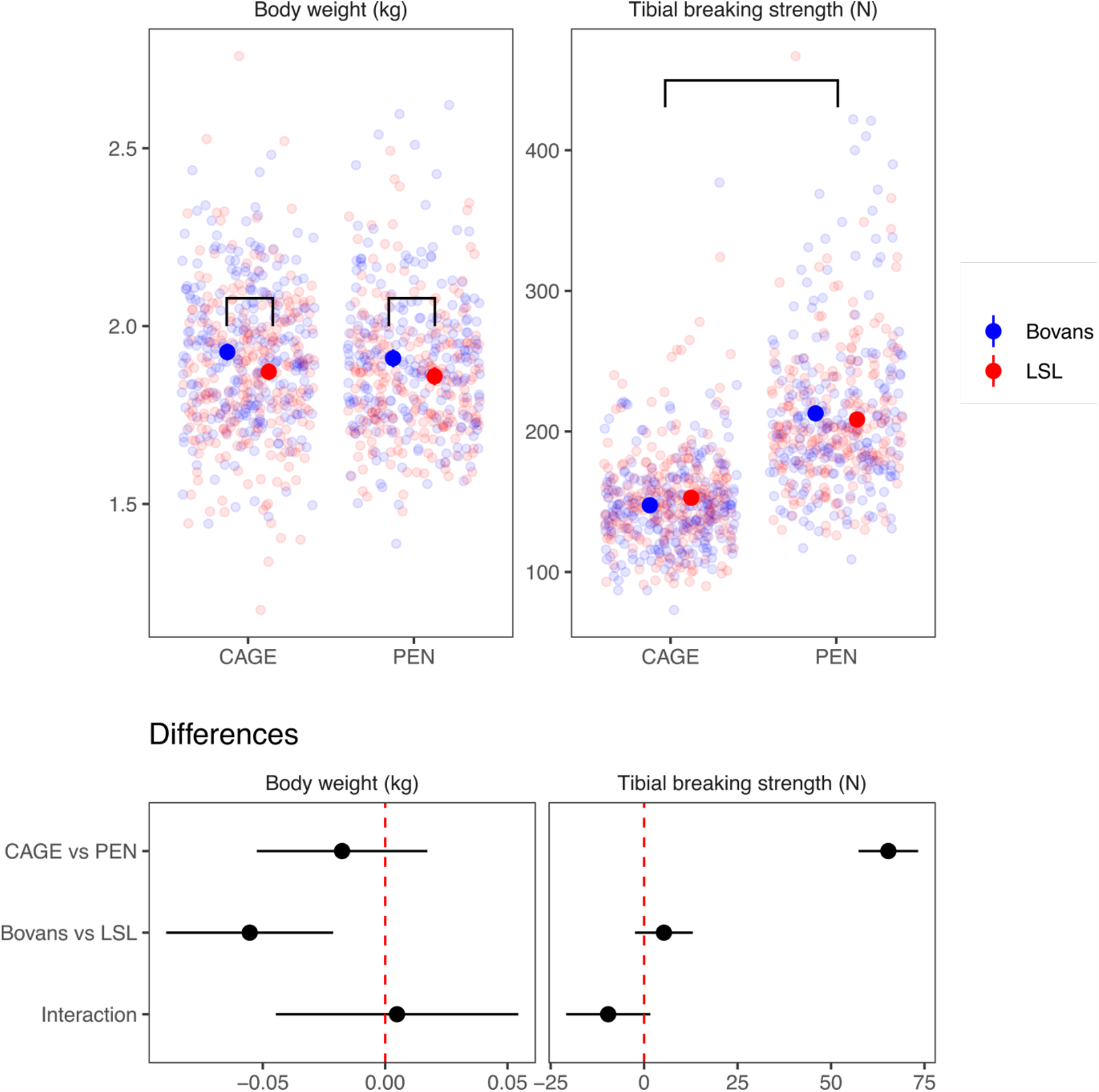
Differences in bone strength between housing systems. Body weight and tibial breaking strength broken down by housing system and crossbred, and estimates of differences between housing systems and crossbreds from a linear model including housing system, breed and an interaction term. The error bars are 95% confidence intervals. The brackets indicate significant differences in body weight between breeds and bone breaking strength between housing systems.

**Figure 2.**
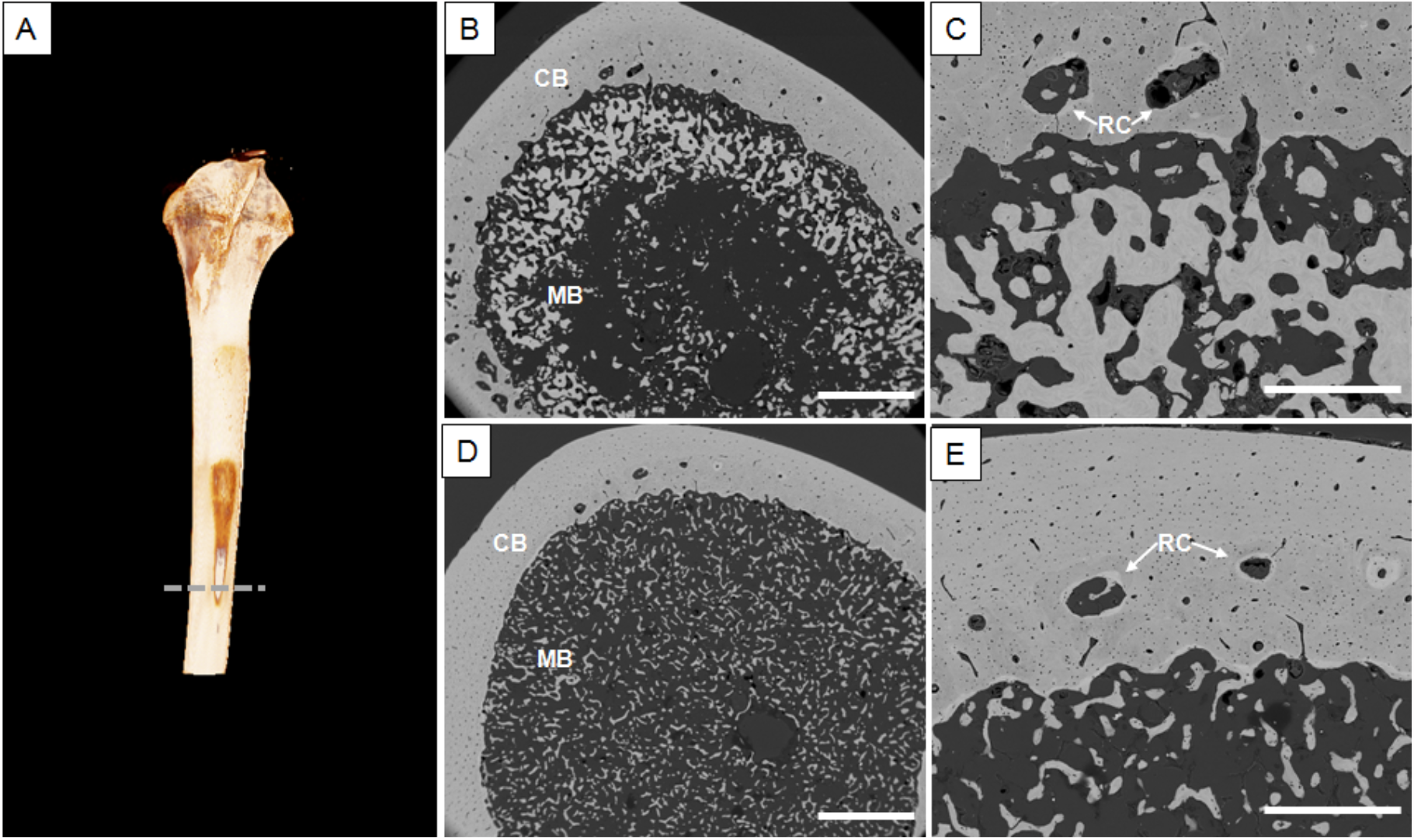
A) 3D image of a tibiae reconstructed from micro-CT. Electron backscattering images of tibia cross-section at mid-shaft from hens of different groups: PEN (B-C) and CAGE (D-E). CB: cortical bone. MB: Medullary bone. RC: resorption center. Scale bar B and D: 1 mm; C and E: 400 µm. Pen birds shows a greater amount of medullary bone particles near the endosteal surface.

Also as expected, there was a positive relationship between body weight and tibial breaking strength in both systems, explaining around 10% of the variance in tibial breaking strength. Figure 3 shows scatterplots of tibial breaking strength and body weight with regression coefficients from a linear model, showing a positive relationship between body weight and bone strength regardless of housing system.

**Figure 3.**
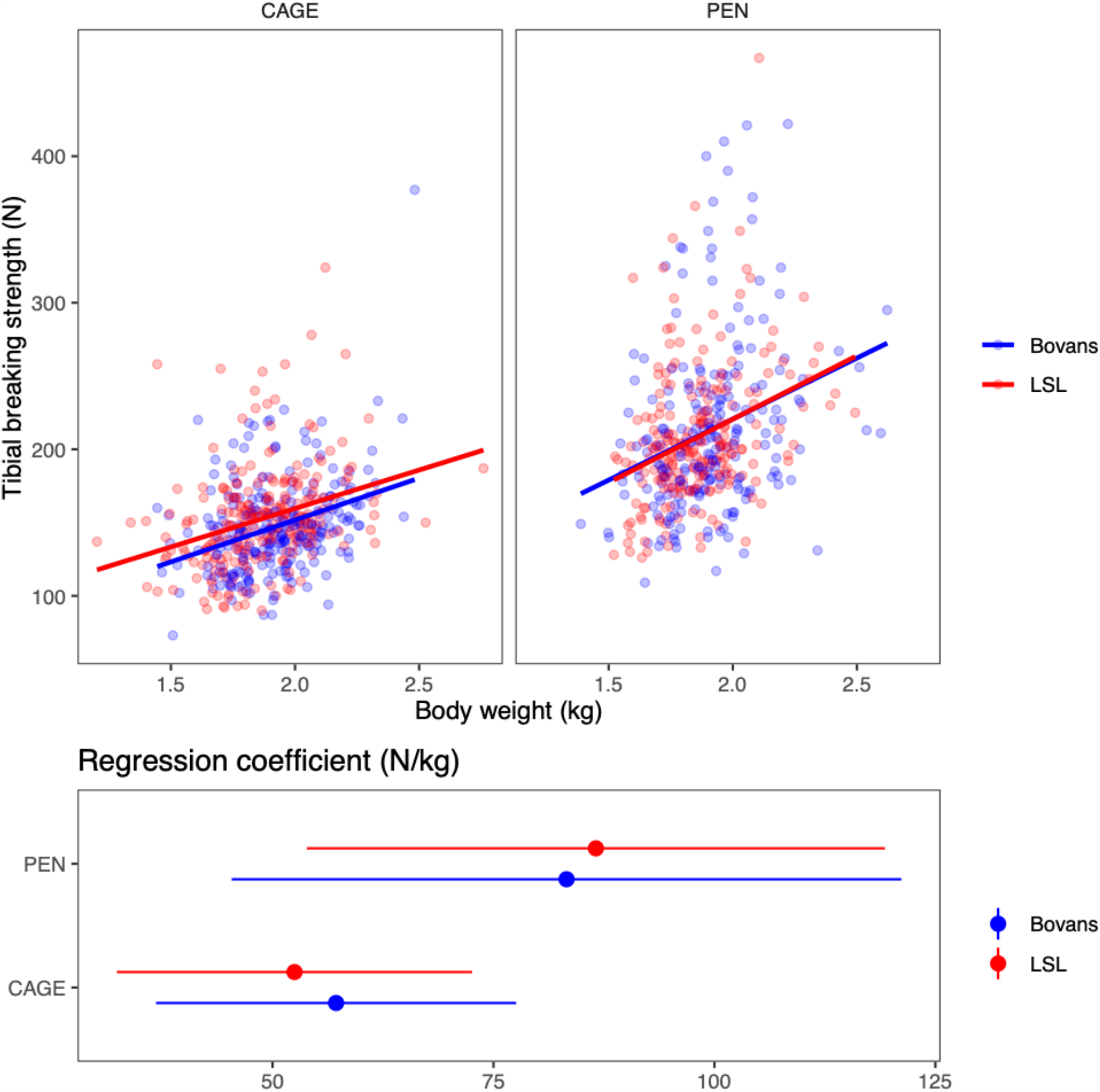
Relationship between bone strength and body weight. Tibial breaking strength and body weight broken down by housing system and breed and regression coefficients from a linear model within breed and housing system. The error bars are 95% confidence intervals.

These differences in bone strength between housing systems were accompanied by differences in bone geometry, mineral content, cortical thickness and bone mineral density (as measured by quantitative computed tomography, QCT) and chemical composition (as measured by thermogravimetric analysis, TGA) between the housing systems. Figure 4 shows heatmaps of the correlations between these bone biomechanical properties, broken down by housing system. Figure 5 shows estimates from a linear model for the first three principal components of the QCT measurements and the main bone composition phenotypes from thermogravimetric measurements (Supplementary Figure 4 shows all variables). Overall, there were differences between the housing systems in most aspects of bone content and composition.

**Figure 4.**
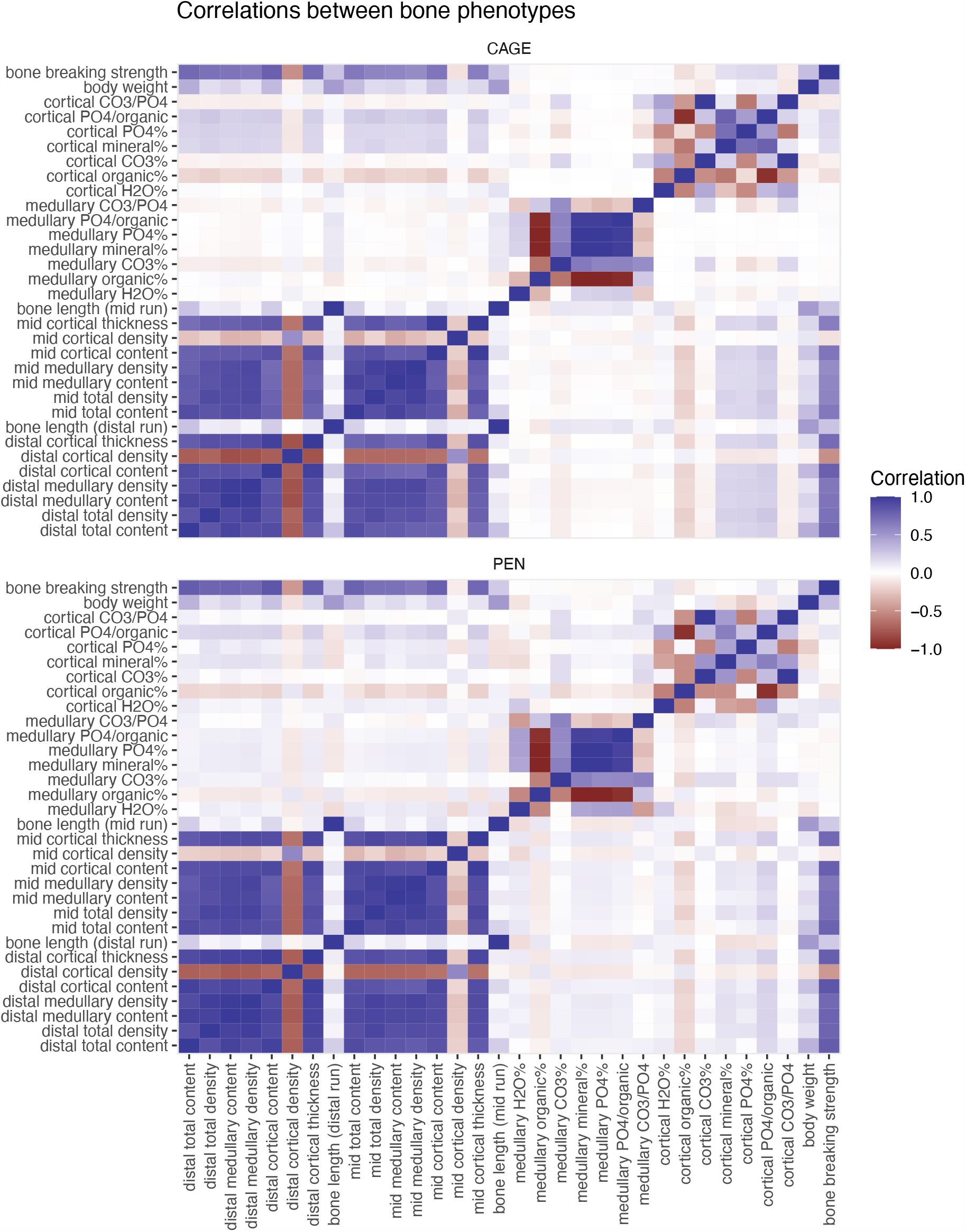
Correlations between bone phenotypes and body weight. The heatmaps show Pearson correlation of body weight, bone breaking strength, density, thickness, content and bone composition traits, separated by housing system.

**Figure 5.**
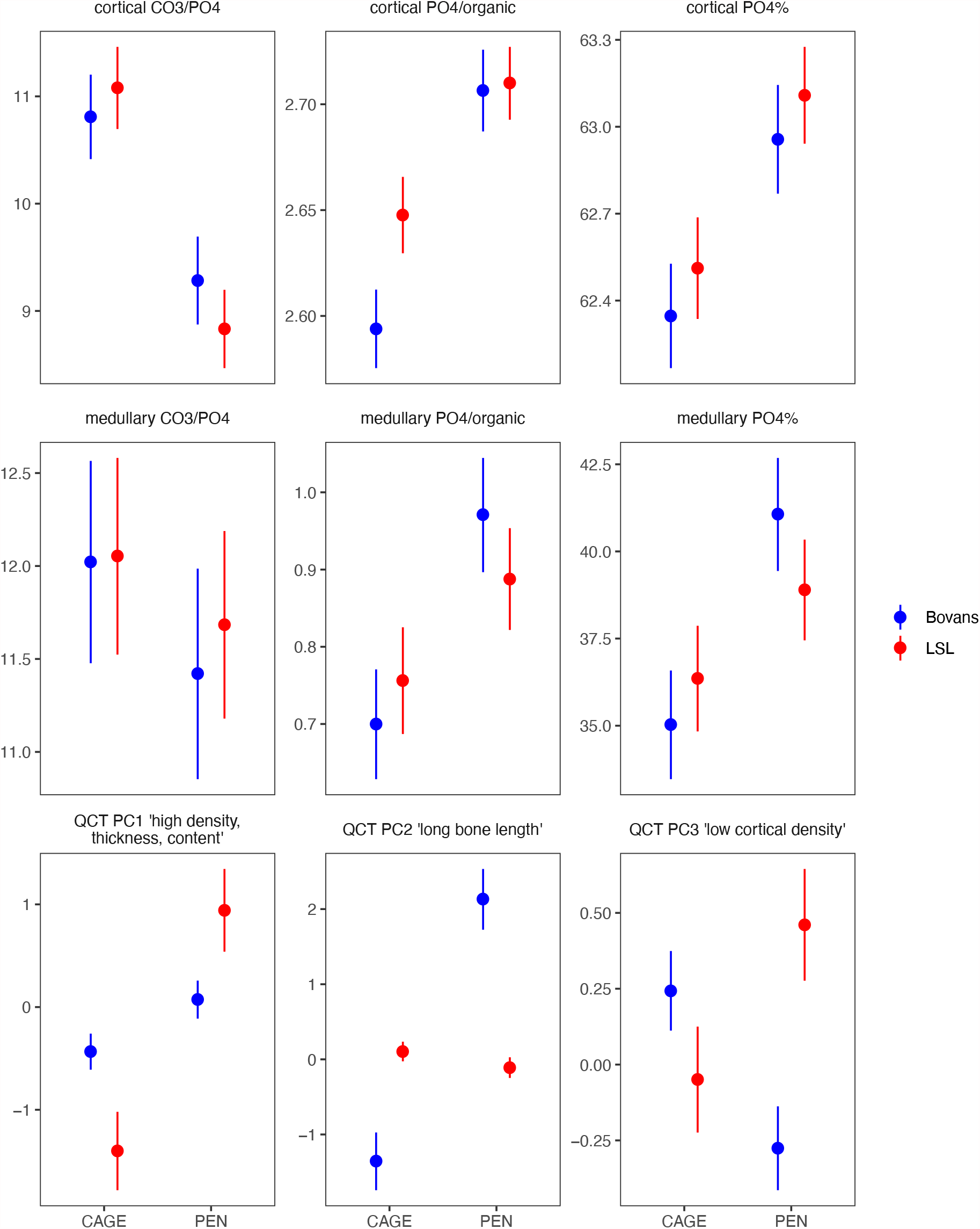
Differences in main bone phenotypes between housing systems. Estimates of means broken down by housing systems and crossbreds from a linear model including housing system, breed and an interaction term, with 95% confidence intervals. All the within breed comparisons between housing systems, except for medullary CO3/PO4, are significant at the p < 0.05 level.

The first QCT principal component, for which cortical density, thickness and bone mineral content had the largest contributions, also show that bone quality was improved in hens housed in a floor pen. Also, the tibia of hens housed in pens had cortical bone with a greater degree of mineralisation and a larger amount of medullary bone than hens housed in furnished cages, as indicated by the PO4/organic and PO4% parameters determined by TGA for both types of bone (Figure 5). Additionally, there were differences in bone chemical composition, such as the amount of carbonate (CO3/PO4) in the cortical bone mineral was significantly lower for hen housed in pens than those housed in furnished cages (Supplementary Figure 4).

### Heritability of bone phenotypes

Bone strength, body weight and tomographic phenotypes had moderate to high genomic heritability. Table 1 shows the estimated genomic heritability and the p-value of a likelihood ratio test for the genomic additive genetic effect. The estimates for bone composition traits (measured by thermogravimetric analysis) were generally lower and most of them were not statistically significant at the p < 0.05 level. It should be noted that this analysis has a smaller sample size than the bone strength and tomographic traits.

**Table 1.**
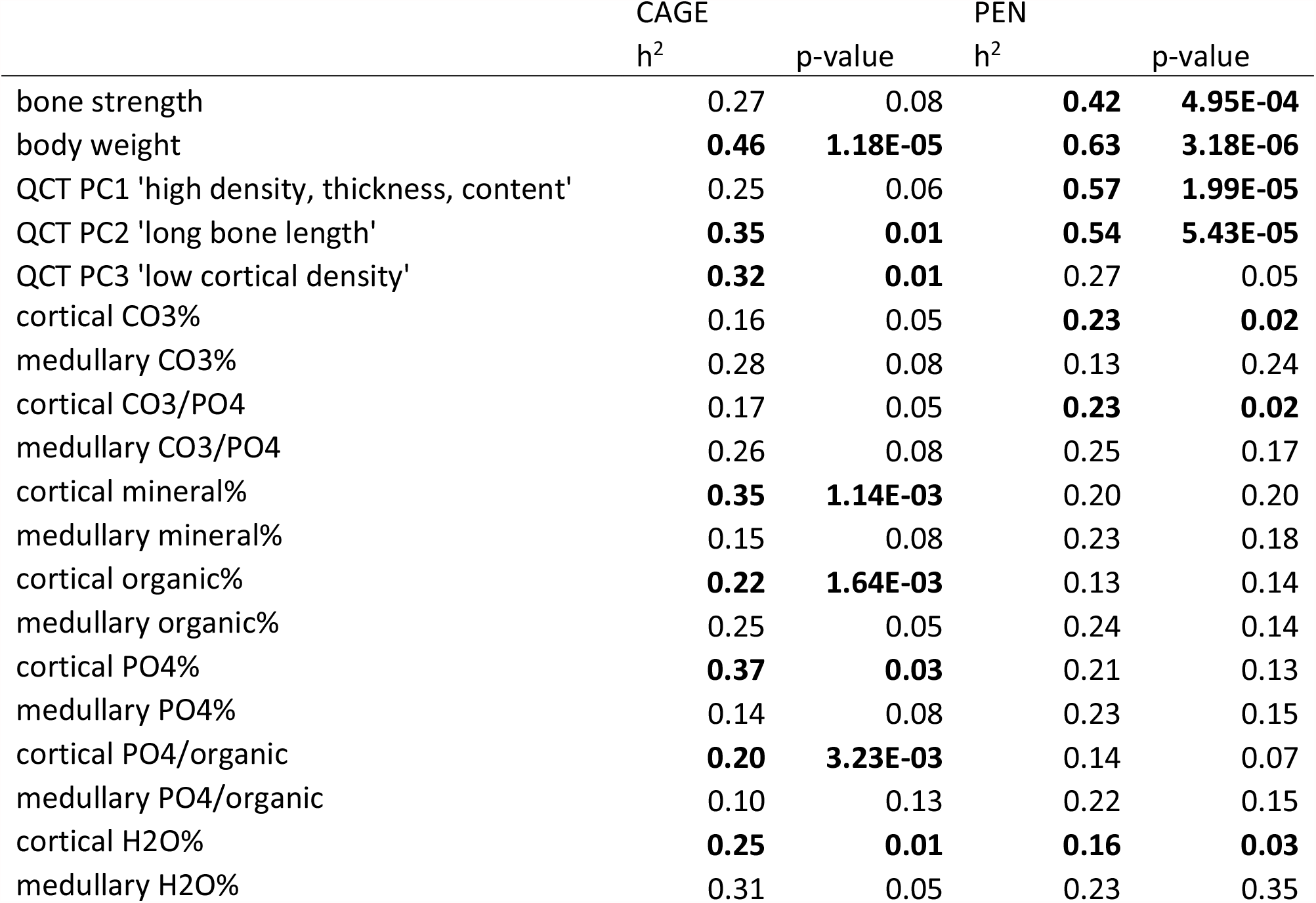
Heritability estimates for bone phenotypes and body mass, separated by cage and pen, with p-values from a likelihood ratio test for the genomic variance component. Bold estimates are significant at the p < 0.05 level.

### Genome-wide association for bone strength and body weight

Genome-scans either combining or independently analysing the two housing systems detected no genome-wide significant loci for bone strength (p < 5 * 10^−8^), but five suggestive loci (p < 10^−4^). Figure 6 shows Manhattan plots of the genome-wide association studies for bone strength, analysing the housing systems jointly and independently. Supplementary figure 6 shows quantile-quantile plots, and Supplementary figure 7 a zoomed-in view of the suggestive loci.

**Figure 6.**
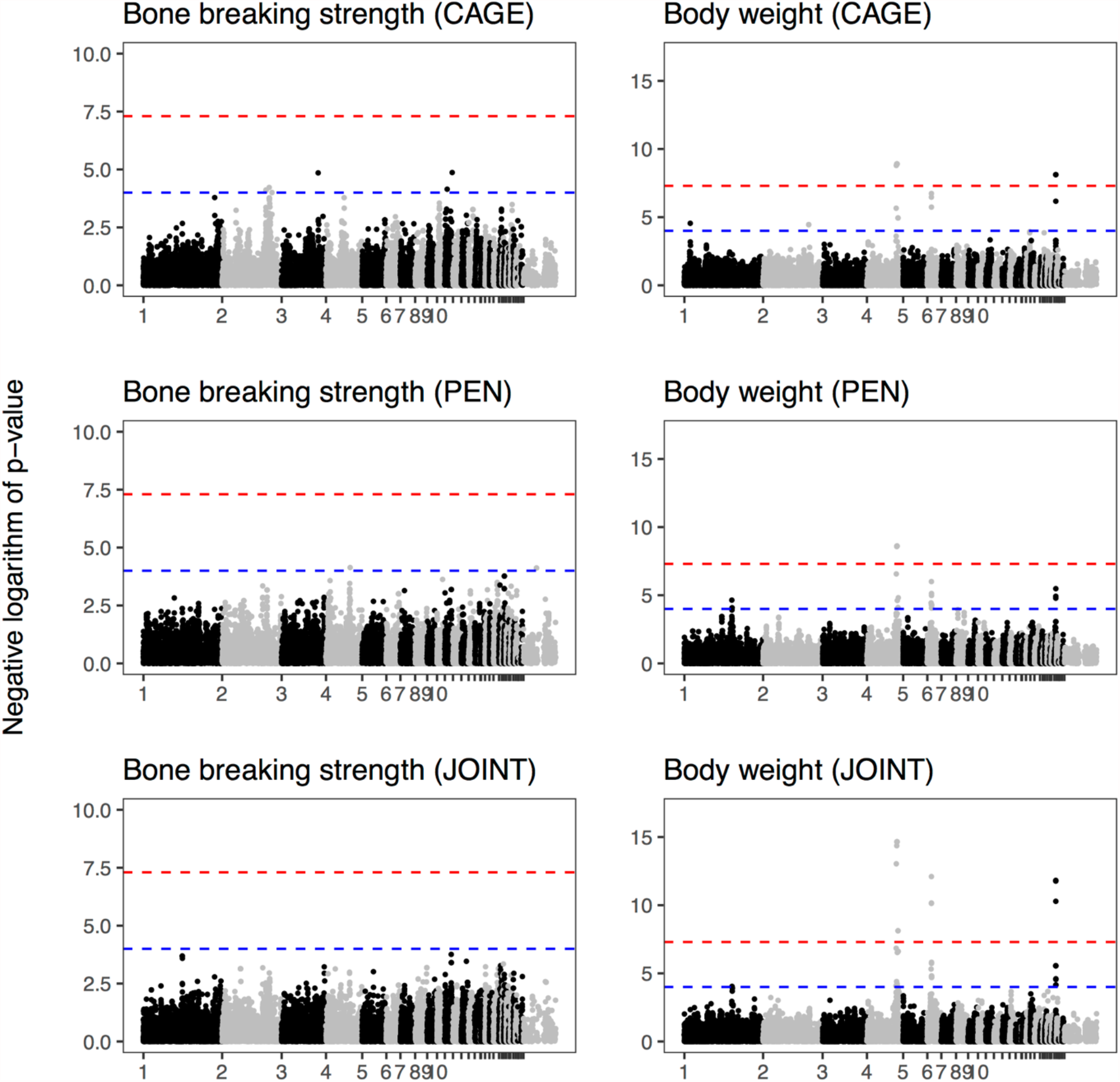
Genome-wide association of bone strength and body weight. Manhattan plots from genome-scans of bone strength and body mass, either separating the housing systems or combining them. Bone strength scans included body mass and crossbred, and in the case of the joint scan also housing system, as fixed effects as well as random effects for housing groups (see Methods). Body weight scans included crossbred, and in the joint scan also housing system, as fixed covariates, as well as random effects for housing group. Chromosome names of the smaller chromosomes have been suppressed for legibility. The dashed red line shows a conventional genome-wide significance threshold of 5 * 10^−8^, and the dashed blue line a suggestive threshold of 10^−4^.

The suggestive associations with bone strength did not overlap previously detected candidate regions for bone strength defined from other populations (Supplementary Table 2). However, there were markers with p < 0.01 in three of these regions, on chromosomes 2, 8 and 23 (Supplementary Table 3).

We detected three significant loci for body weight on chromosomes 4, 6 and 27 that were either significant (p < 5 * 10^−8^) or suggestive (p < 10^−4^) in both the joint and separate scans. Supplementary Figure 8 shows a zoomed in view of the three body weight loci. Table 2 shows the locations of significant associations.

Because the chromosome 4 locus contains multiple significant markers spread over a region of several megabasepairs, we performed a conditional scan that included the most significant marker in the region as a covariate (Supplementary Figure 9). Controlling for the most significant marker abolished the significant association throughout the whole region, meaning that we have no clear evidence of multiple linked loci in the region.

**Table 2.** Overview of significant regions from genome-wide association scans.

| Trait | Chromosome | Lead SNP position | Lead SNP p-value |
| --- | --- | --- | --- |
| QCT PC2 'long bone length' (PEN) | 2 | 55321235 | 4.87E-08 |
| QCT PC2 'long bone length' (JOINT) | 2 | 55833673 | 2.98E-09 |
| QCT PC3 'low cortical density' (CAGE) | 4 | 73994772 | 2.49E-09 |
| body weight (CAGE) | 4 | 75151189 | 1.24E-09 |
| QCT PC2 'long bone length' (CAGE) | 4 | 75151189 | 1.51E-08 |
| body weight (PEN) | 4 | 75748329 | 2.45E-09 |
| body weight (JOINT) | 4 | 75748329 | 2.22E-15 |
| QCT PC2 'long bone length' (JOINT) | 4 | 75748329 | 3.02E-11 |
| body weight (JOINT) | 6 | 11477631 | 7.91E-13 |
| QCT PC2 'long bone length' (CAGE) | 6 | 11477631 | 8.40E-11 |
| QCT PC2 'long bone length' (JOINT) | 6 | 11477631 | 2.26E-13 |
| body weight (CAGE) | 27 | 6070932 | 7.60E-09 |
| body weight (JOINT) | 27 | 6087051 | 1.52E-12 |
| QCT PC2 'long bone length' (PEN) | 27 | 6087051 | 9.52E-09 |
| QCT PC2 'long bone length' (JOINT) | 27 | 6087051 | 6.62E-11 |

### Genetic differences between housing systems

There was no overlap between the suggestive loci for bone strength in the two housing systems. Figure 7 compares the p-values and estimated marker effects, using all markers with p < 10^−3^ between the floor pen and furnished cage systems. For comparison, we also show the same scatterplots for the body weight scan, where the loci overlap between housing systems.

**Figure 7.**
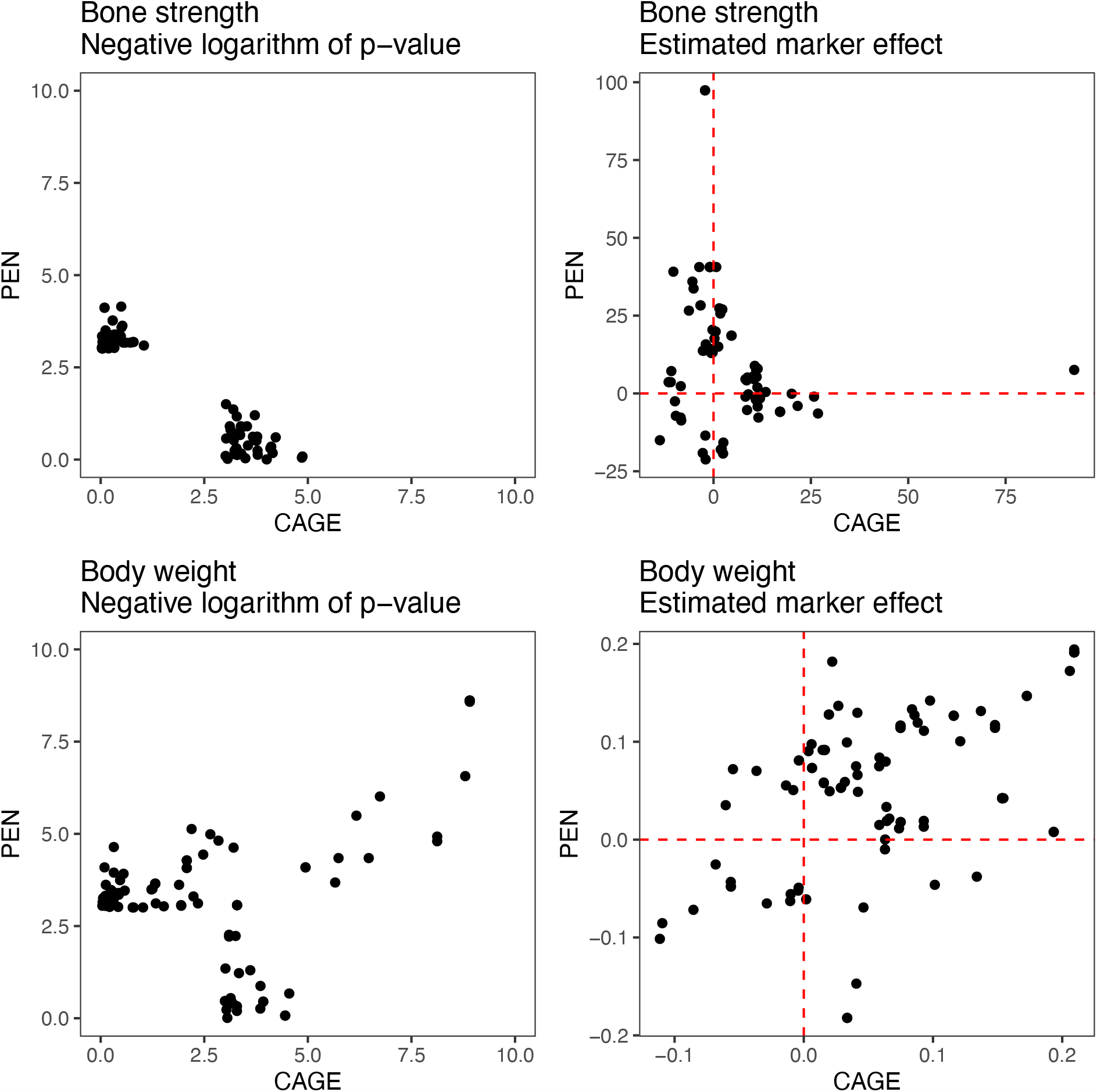
Comparison of genetic associations between housing systems. Scatterplots compare the p-values and estimated marker effects of markers with p < 10^−3^ either in furnished cages or in floor pens.

Genetic correlation estimates between housing systems were too imprecise to be useful. We used a bivariate model with the genomic relationship matrix to estimate genomic heritability and correlation between housing systems. Supplementary Table 4 shows the estimated genetic correlations and heritabilities from this model.

### Genome-wide association of bone mineral density and bone composition

Genome-scans for the ten bone mineral density and bone composition phenotypes that had significant heritability detected five more significant associations and 50 suggestive associations. Figure 8 show Manhattan plots of QCT phenotypes, which had significant associations. Supplementary Figures 11 and 12 show Manhattan plots for the genome scans for thermogravimetric phenotypes that had suggestive associations. Table 2 and Supplementary Table 5 summarise the location of significant and suggestive regions, respectively. There were four significant associations for the second principal component of QCT phenotypes, reflecting bone length (Figure 8). Three of them coincided with the body weight loci on chromosomes 4, 6 and 27. There was also another significant association on chromosome 2. The third principal component, reflecting cortical thickness and content, had one significant association, coinciding with the body weight association on chromosome 4.

**Figure 8.**
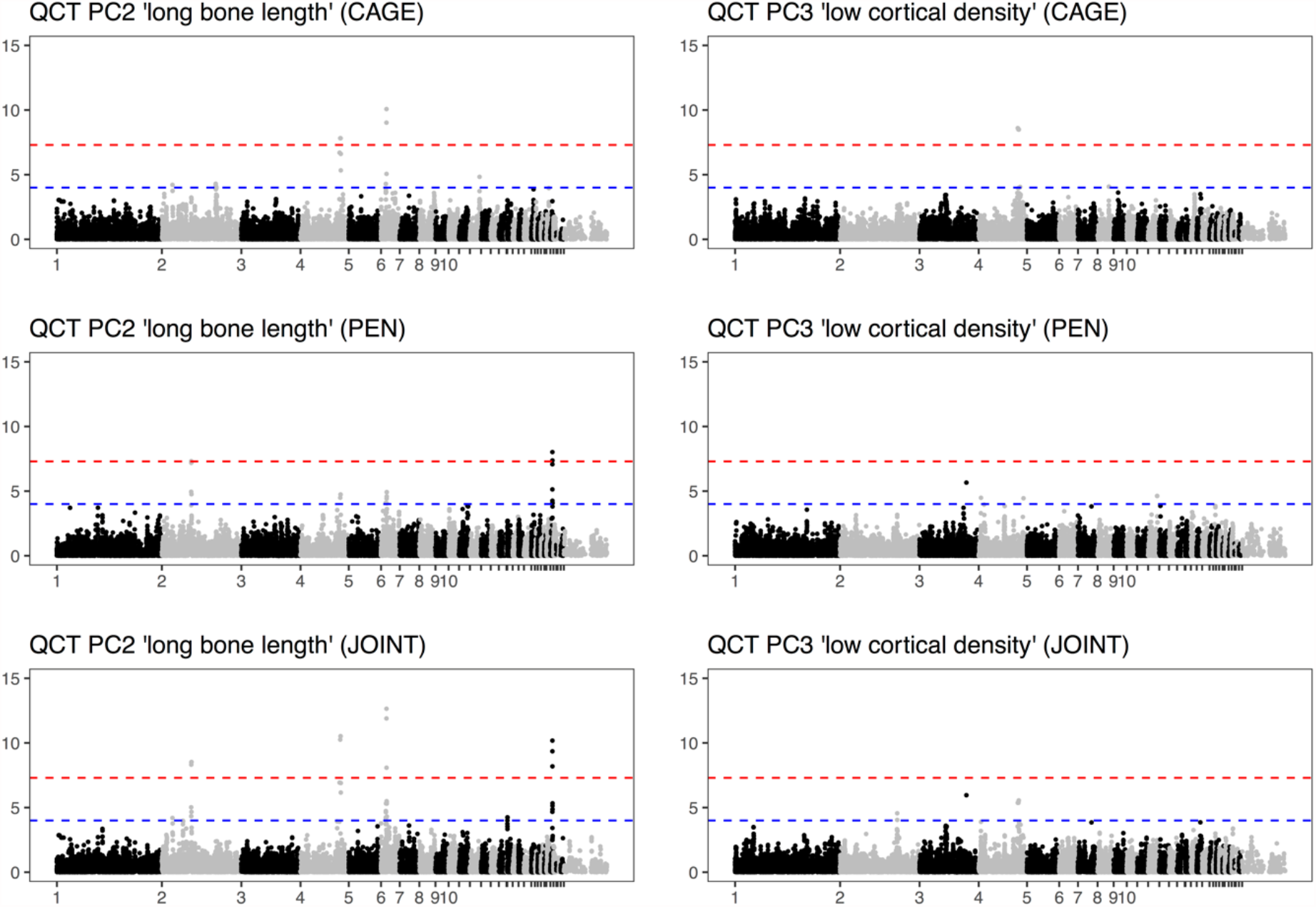
Genome-wide association of the second and third principal components of QCT phenotypes, which had significant heritability both housing systems. Genome scans included body mass and crossbred, and in the case of the joint scan also housing system, as fixed effects as well as random effects for housing groups (see Methods). Chromosome names of the smaller chromosomes have been suppressed for legibility. The dashed red line shows a conventional genome-wide significance threshold of 5 * 10^−8^, and the dashed blue line a suggestive threshold of 10^−4^.

## Discussion

In this paper, we found that bone strength in commercial crossbred laying hens is highly polygenic and potentially exhibits gene-by-environment interactions between housing systems that allow different amounts of exercise. We detected no genome-wide significant loci for bone strength, and the suggestive loci were different between the two environments. In contrast, we detected three significant body weight loci shared between environments and coincided with significant loci for bone length. This leads to three topics for discussion:

1. differences in bone strength, content and composition between floor pen and furnished cage housing systems;
2. evidence for gene-by-environment interaction between housing systems;
3. candidate genes underlying loci for body weight and bone length.

### The effect of housing system on bone strength, content and composition

Our detailed bone phenotyping revealed several differences in bone strength, content and chemical composition between hens housed in furnished cages and hens housed in floor pens. The environmental difference between housing systems causes a quantitative increase in bone strength accompanied by increased bone formation and mineralisation and in the floor pen system, where the hens are able to exercise more.

In addition to greater bone strength, the QCT results show the principal component containing predominately cortical density, cortical thickness and bone mineral content being improved in a pen environment. Previous results also demonstrated increased bone cortical thickness, a lower bone cortical porosity, a larger amount of medullary bone and overall a greater total bone mass as factors contributing to the greater strength seen in hens housed in aviary systems that also allowed for greater mobility (Fleming et al., 2006; Rodriguez-Navarro et al., 2018; Shipov et al., 2010). Also, analysis by thermogravimetry show that hens housed in floor pens have a higher degree of bone mineralization. The main traits describing the amount of bone mineralization of cortical and medullary bone (PO4/organic and PO4%) were greater in hens housed in floor pens than in furnished cages. This is consistent with previous results, as the greater opportunity for physical exercise stimulates bone formation and increases mineralisation of the medullary cavity (Rodriguez-Navarro et al., 2018; Shipov et al., 2010).

On the other hand, hens in floor pens had bone with a greater degree of mineralisation and a higher carbonate/phosphate ratio than hens housed in cages. A greater degree of mineralization and lower carbonate/phosphate ratio is indicative of an increased bone maturity and lower turnover rates reflecting a decreased amount of remodelling of established bone in hens in floor pens. In contrast, Rodriguez-Navarro et al. (2018) found that hens with increased mobility had cortical bone with lower degree of mineralisation and higher carbonate/phosphate ratio, suggesting a higher amount of bone remodelling. Thus, it seems the effect of exercise on bone remodelling and maturation also depends on other factors, such as age or other environmental variables.

This discrepancy in the response of bone to physical activity might be explained by aging effects, if a higher metabolic activity in pen-housed chickens at an earlier age coincides with, or even causes, a lower metabolic activity at a later age. The hens in Rodriguez-Navarro et al. (2018) were 56 weeks old at sampling, while the hens in this study were 100 weeks old; the differences in bone strength and geometry might have been established at an earlier ages. Bone metabolism is a dynamic process where what happened earlier in life matters. For example, bone quality is negatively genetically correlated with age at first egg, suggesting that early sexual maturation causes worse bone quality later in life (Dunn et al., 2021). Similarly, whether pullets are reared in cages or in aviaries, allowing for more movement, has long-term effects on bone properties later in life (Casey-Trott et al., 2017). This suggests that longitudinal studies of bone mineralisation and remodelling in layer hens are warranted.

As we have observed before, the medullary bone shows more pronounced effects than cortical bone. Thus, it appears that medullary bone responds to exercise even at older age, despite contributing less to bone strength than cortical bone. This is in accordance with previous results: Medullary bone composition had significant heritabilities in white and brown egg layers (Dunn et al., 2021), and medullary bone has showed increased PO_4_/amide levels in response to exercise (Rodriguez-Navarro et al., 2018; Shipov et al., 2010). Previous studies also suggested that the amount of medullary bone was increased by the selection for better bone quality and by increased physical activity in aviary systems (Fleming et al., 2006). Medullary bone was clearly more mineralised in both breeds when housed in pens. In this study, there is little apparent correlation between medullary bone and bone strength, but other studies have found association between medullary mineralisation and bone strength (Alfonso-Carrillo et al., 2021; Rodriguez-Navarro et al., 2018). Thus, variation in medullary bone is an important contributor to variability in bone mineral content and mechanical properties, both in terms of genetic variation and response to exercise.

### The evidence for gene-by-environment interaction between housing systems

Genome-wide association scans of bone strength gave completely different results between hens housed in furnished cages and hens housed in floor pens, suggesting that the genetic basis of bone strength may be different in the two housing systems. There were no suggestive associations in common between the two housing systems, and little concordance between estimated marker effects. In combination with evidence for differences in bone content and composition between housing systems, we hypothesise that this difference is due to gene-by-environment interaction. That is, the genetic architectures of bone strength in a furnished cage and in a floor pen are different, likely because these environments put such different pressures on bone development and homeostasis. Therefore, the genes involved in bone turnover in response to loading may be substantially different to those involved in contributing to variance where loading is less. On the contrary, the genome-wide association results for body weight were consistent between the housing systems. This similarity suggests that the genetic variants that affect growth, at least at the three loci detected in this study, do not interact with the housing system. At the same time, there was little difference in body weight between hens in the two housing systems.

Low power to detect associations is unlikely to explain this pattern of gene-by-environment interaction. A previous genome-wide association study in a homogenous group of 750 pure line hens detected several strong associations for bone strength (Raymond et al., 2018). The pure line hens were from the Lohmann breeding program, and therefore closely related to one of the crossbreds used in the current study. Thus, a study of this size would likely be powered to detect loci for bone mineral density in the absence of gene-by-environment interaction, as it is with loci for body weight that are shared between environments. Therefore, the lack of shared associations for bone strength between housing systems are unlikely to be explained by low power to detect them. If the previously known loci had similar effects in both environments, we should be able to detect them. For context, estimated additive genetic effects detected by Raymond et al., (2018) range from 11 to 33 N, which is comparable to the additive effects estimated within housing system in this study (ranging from 7 to 21 N). These effects can be compared to the average difference between housing systems, which is 65 N.

### Candidate genes for body weight and bone length

The body weight loci on chromosomes 4, 6, and 27 overlap loci reported in several previous genetic mapping studies. The regions overlap several compelling candidate genes for body weight in chickens, which is also reflected in enrichment of body weight and feed conversion associations from Chicken QTLdb (Supplementary figure 13). This includes studies within laying hen populations where the same region on chromosome 4 was seen to also have pleiotropic effects on a wide range of traits including egg quality traits (Wolc et al., 2014).

Two different loci for body weight overlapping our chromosome 4 locus have been fine mapped down to regions of one or a few candidate genes. A series of genetic mapping studies (Lyu et al., 2018, 2017; Nassar et al., 2015) detected and progressively fine-mapped a region containing 15 genes, including *Ligand dependent nuclear receptor corepressor like* (*LCORL; ENSGALG00000014421*) and *Condensin complex subunit 3* (*NCAPG*; *ENSGALG00000014425*). This locus is also associated with body size traits in humans (Weedon et al., 2008), cattle (Bouwman et al., 2018) and horses (Makvandi-Nejad et al., 2012). The other locus was detected by (Sewalem et al., 2002) and fine mapped to *Cholecystokinin receptor type A* (*CCKAR*; *ENSGALG00000030801*), and was shown to alter the expression of the *CCKAR* gene and the physiological response of the animals to its ligand CCK (Dunn et al., 2013). The associated region found in this study overlaps both of these regions. One or both of them might contribute to the association; due to linkage disequilibrium, we cannot tell them apart. We confirmed this by a conditional genome-wide association scan, where adding the lead SNP as a covariate abolished the association signal throughout the region. This suggests that linkage disequilibrium throughout the region prevents us from genetically dissecting it further in this population. This region appears to be a hotspot of genetic effects on body weight, or perhaps more correctly stature, across a large range of animals, with pleiotropic effects on other traits.

The two most significant associations on chromosome 27 fall in the *insulin-like growth factor 2 mRNA binding protein 1* gene (*IGF2BP1*; *ENSGALG00000041204*). *IGF2BP1* is known to be expressed in developing limbs and has been shown to alter the length of chick long bones (Fisher et al., 2005) which could ultimately affect stature. The *IGF2BP1* locus has been highlighted previously in a GWAS study in a population of laying type chicken which included white leghorns genetics and affected a range of carcase traits including feet weight with effects up to 4.78% of the variance (Ma et al., 2019). The study also demonstrated the region between *CCKAR* and *NACPG* as important for carcase traits as in this study. Expression of *IGF2BP1* is also associated with adipogenesis in chickens (Chen et al., 2019). This association is also close to bone candidate gene *sclerotin (SOST; ENSGALG00000009929)*, located about 150 kbp way. Sclerotin is a negative regulator of bone formation that is expressed in osteocytes (van Bezooijen et al., 2005); loss-of-function mutations in humans cause bone overgrowth (sclerosteosis). Guo et al. (2017) report an association with femoral bone mineral content and femoral weight in this region, highlighting *SOST* as a candidate gene. For femoral weight on their lead SNP occurs close to *IGF2BP1*, while their lead SNP for bone mineral content is closest to *SOST*.

We detected significant loci associated with bone length coinciding with the major body weight loci, despite including body weight as a covariate in the bone length genome scan. This may be an artefact of a non-linear relationship between body weight and bone length, or a genuinely pleiotropic effect on bone length. However, there was one association for bone length independent of body weight on chromosome 2. The closest gene was *succinyl-CoA:glutarate-CoA transferase* (*SUGCT; ENSGALG00000031758*). This gene encodes a mitochondrial enzyme that is associated with glutaric aciduria in humans, but appears to have no known connection to bone or to body size traits.

## Conclusion

The current study yet again establishes the positive effects of systems that allow greater movement of laying hens on bone quality, and that these beneficial effects can also be seen in old hens (100 weeks of age). If the unintended consequences of increased collisions in such systems can be reduced by improved design, then the combination of environment, nutrition and genetics, taking in to account what we have learned in this study about environment interactions, then the risk of fracture in laying hens could be minimised. Knowledge acquired in this study could help in moving to selection strategies aimed to reduce the incidence of bone damage in laying hens in systems that allow greater mobility. This might include the use of whole genome selection strategies, even if individual loci that explained large amounts of variance were not detected for bone quality. This could allow phenotypes gathered in extensively housed hens be applied to pedigree hens, which may need to be selected in a cage environment for egg laying performance. This could conceivably be achieved by genomic selection or by sib selection.

## Supporting information

Supplementary Data 1: GWAS summary statistics

Supplementary Table 1: Sample sizes

Supplementary Table 2: candidate regions from previous studies

Supplementary Table 3: candidate association results

Supplementary Table 4: Genomic correlations

Supplementary Table 5: GWAS summary statistics from suggestive regions

## Acknowledgements

The work was supported by Formas – a Swedish Research Council for Sustainable Development projects 2012-1342, 2014-01840 (ERA-NET), and 2016-01386; by the Biotechnology and Biological Sciences Research Council; UK grant ‘Better Bones’ BB/M028291/1 (ERA-NET); and Investigación y Tecnología Agraria y Alimentaria 291815.

## Supplementary figures

**Supplementary Figure 1.**
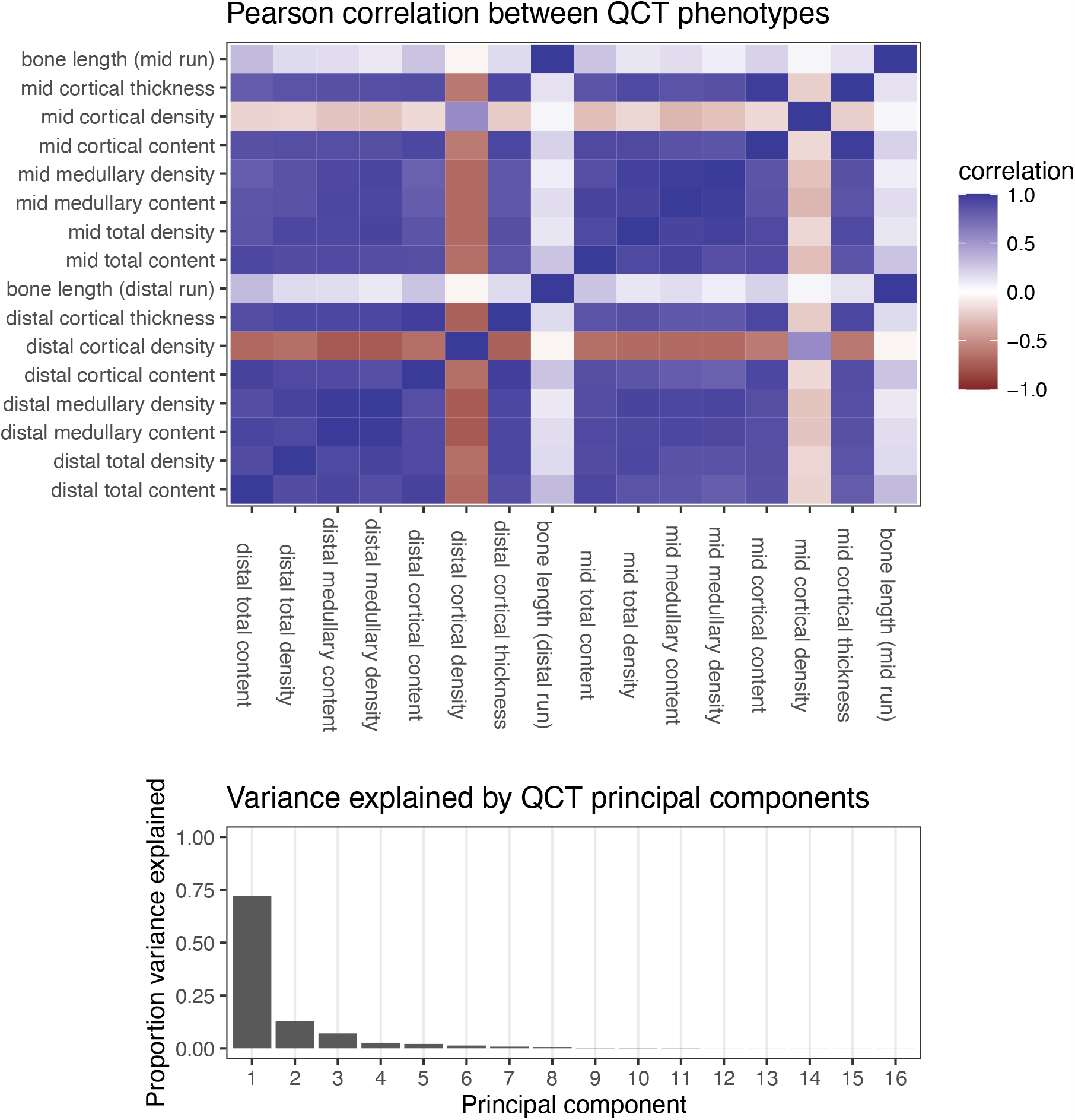
Correlation heatmap and variance explained by principal components of QCT phentoypes.

**Supplementary Figure 2.**
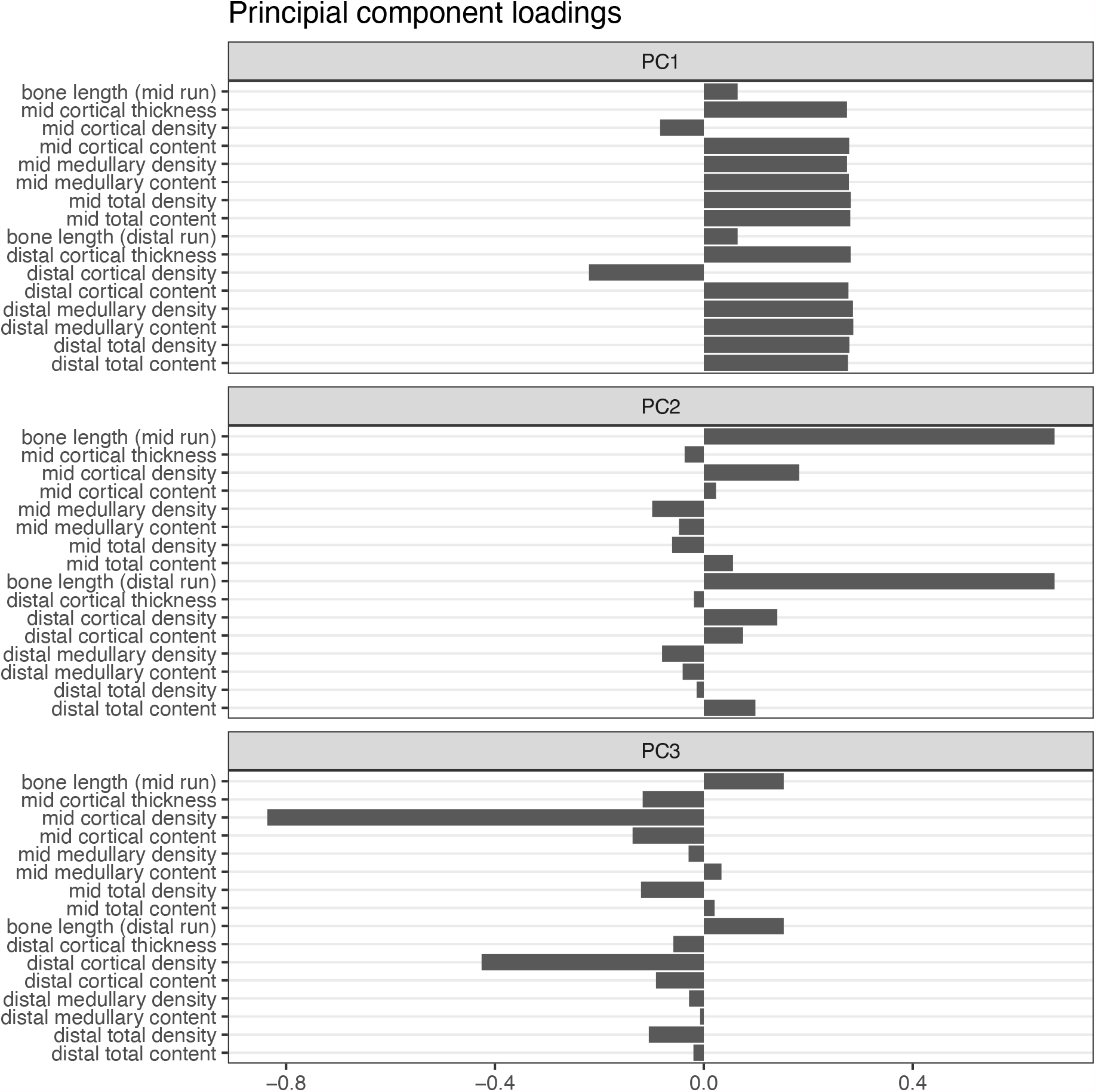
Loadings on the first three principal components of the QCT phenotypes, showing how the first captures most density and content variables, the second tibial bone length, and the third cortical density.

**Supplementary Figure 3.**
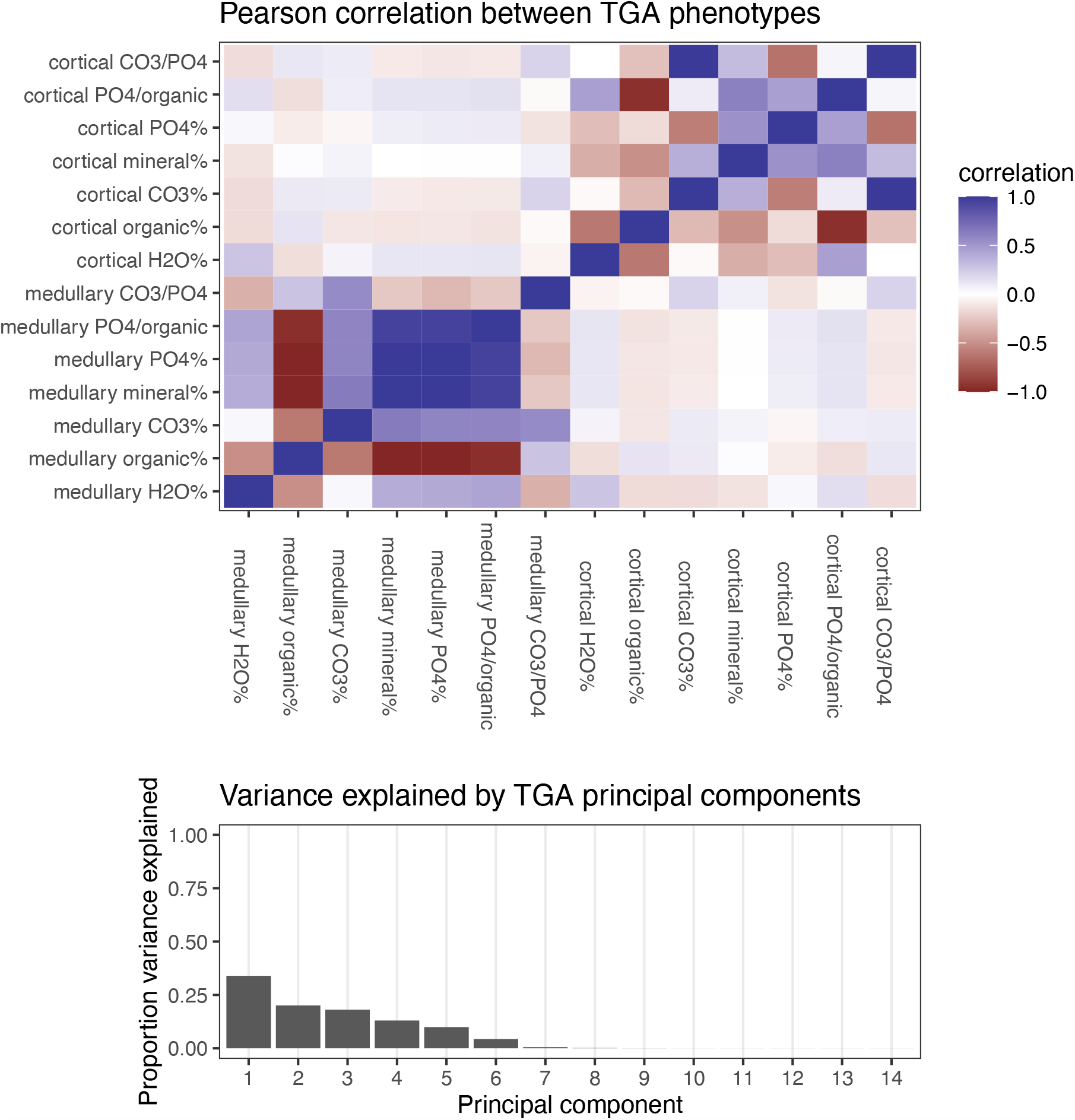
Correlation heatmap and variance explained by principal components of TGA phentoypes.

**Supplementary Figure 4.**
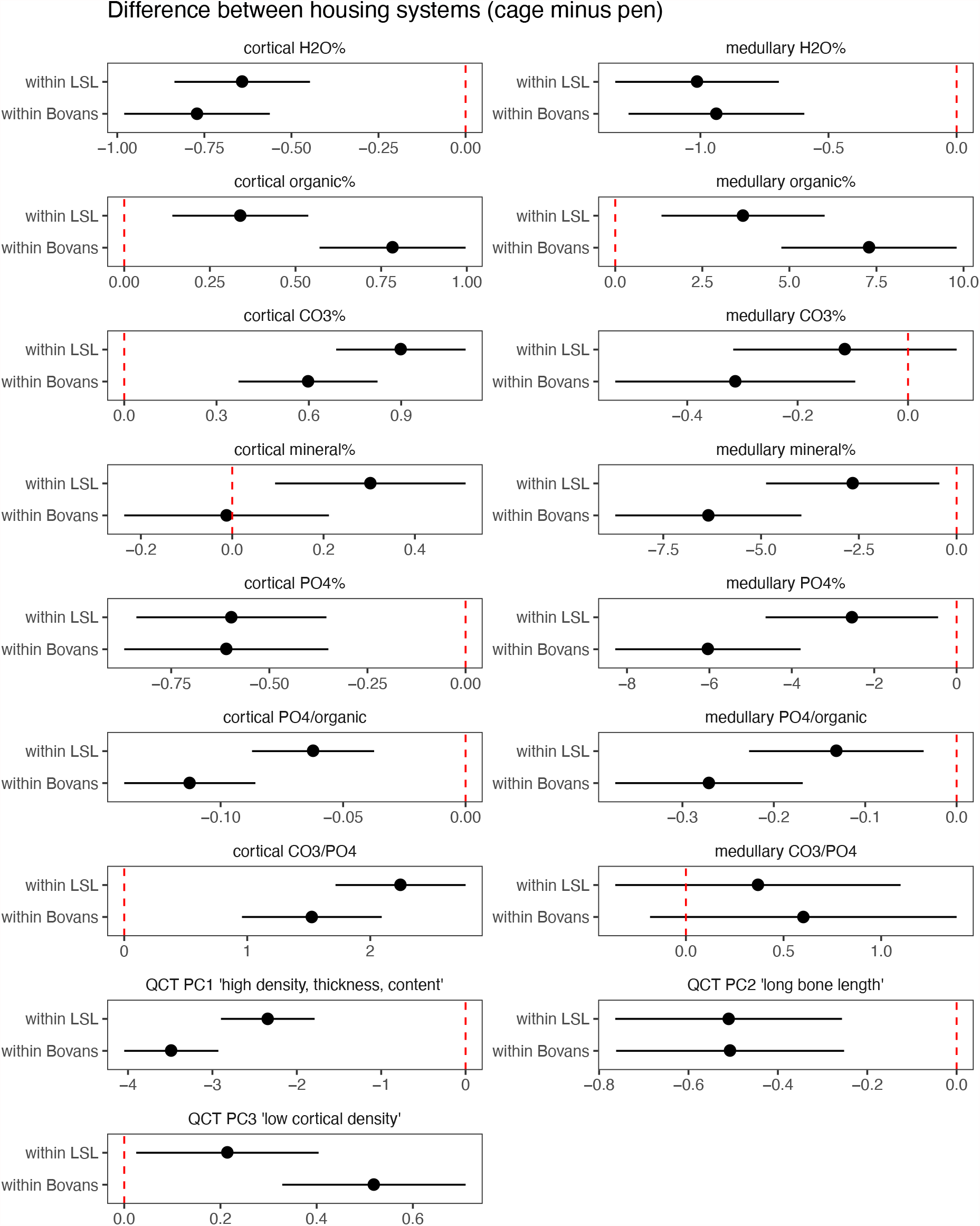
Differences in bone phenotypes between housing systems. Estimates of differences between housing systems and crossbreds from a linear model including housing system, breed and an interaction term. Differences are expressed a linear contrast between housing systems (cage minus pen) within the two crossbreds (LSL and Bovans). Thus, positive values mean that trait values are higher, on average, in furnished cages than in floor pens, and vice versa. The red dashed line indicates zero; intervals that do not overlap this line are significantly different from zero.

**Supplementary Figure 5.**
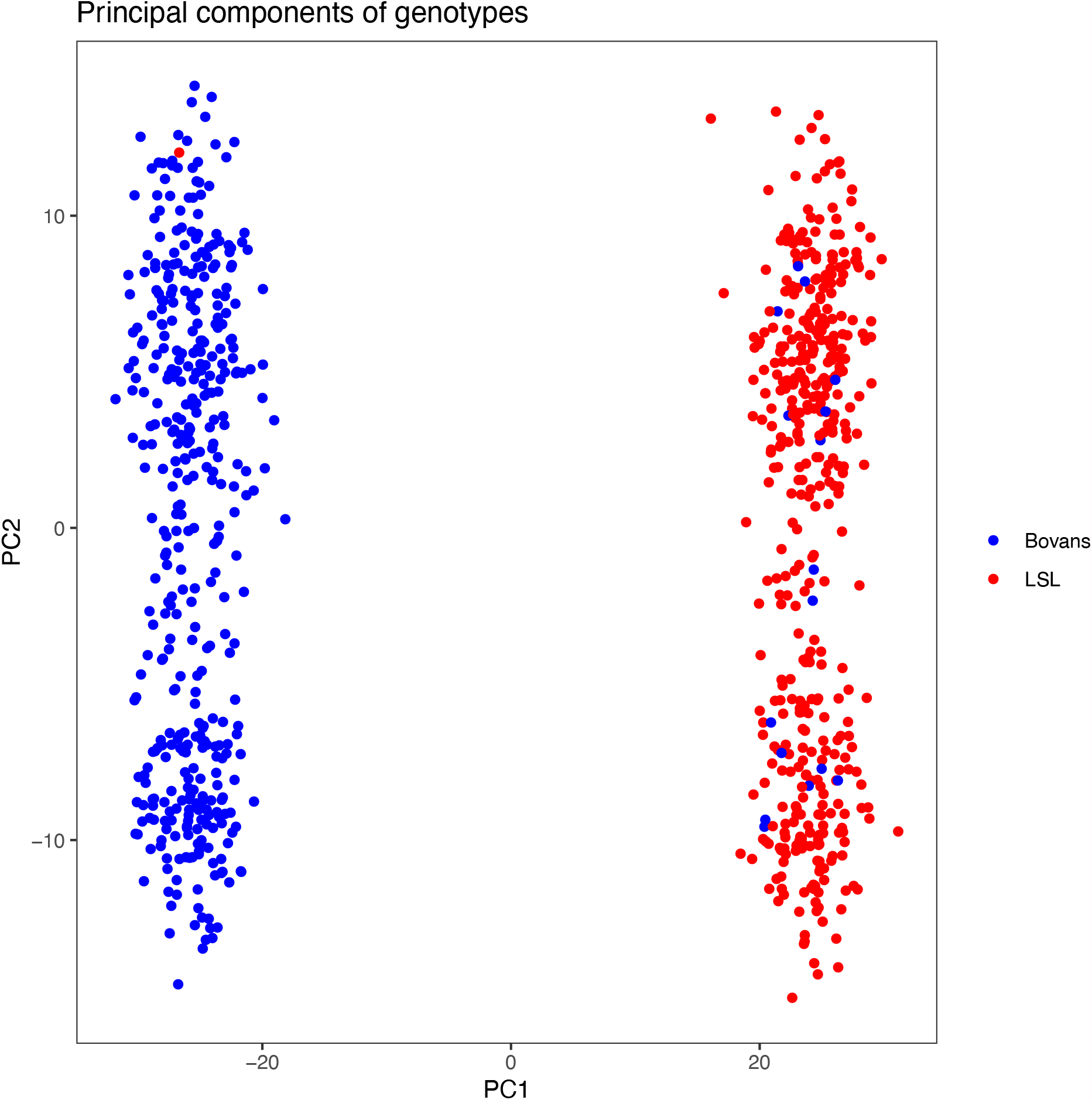
The first principal component separates the two crossbreds. Scatterplot of the first two principal components of the genotypes, coloured by the crossbred. 19 individuals appeared to be recorded as the wrong crossbred based on the position the plot, and were excluded.

**Supplementary Figure 6.**
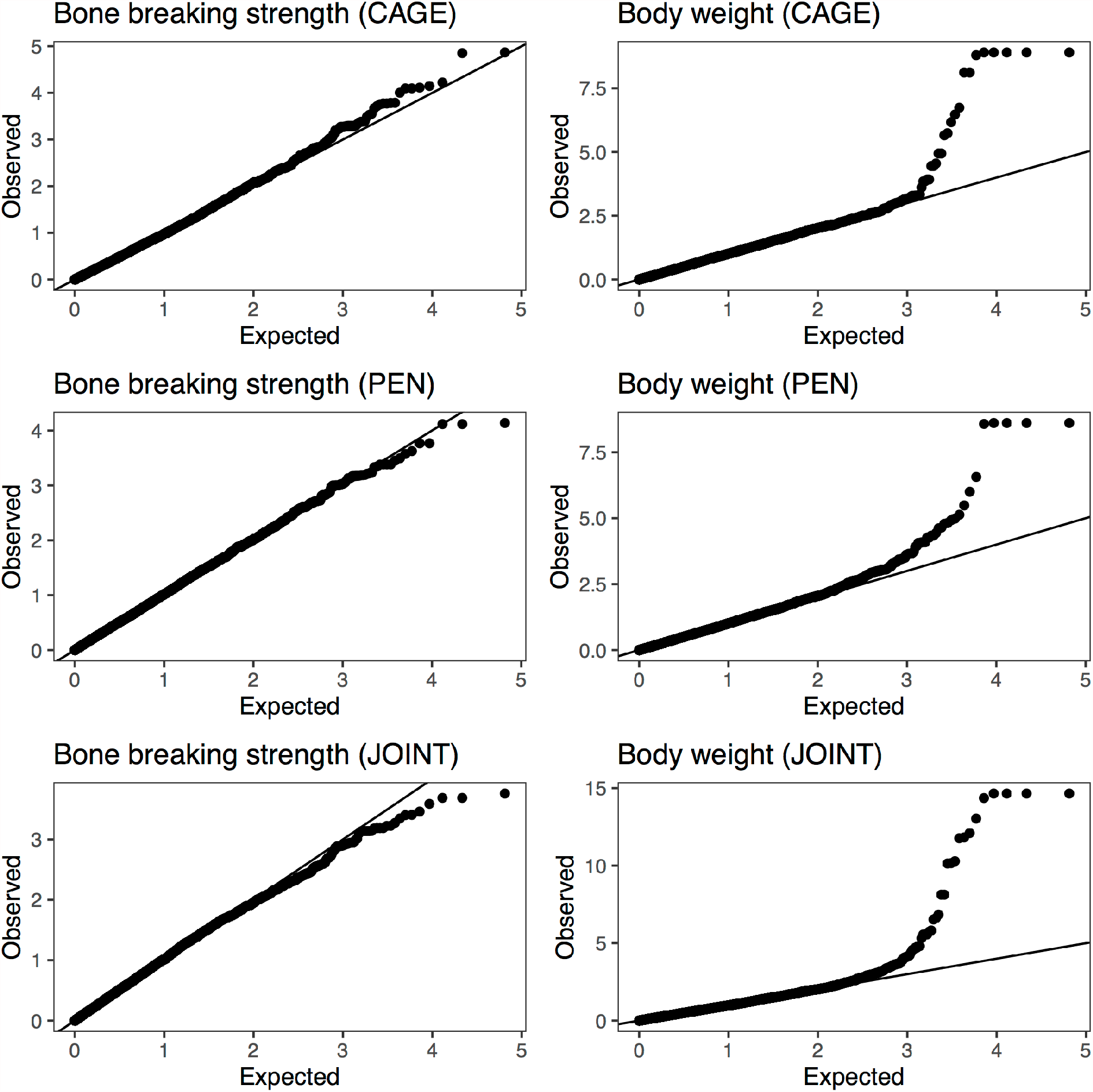
Quantile—quantile plots of genome scans for bone breaking strength and body weight.

**Supplementary Figure 7.**
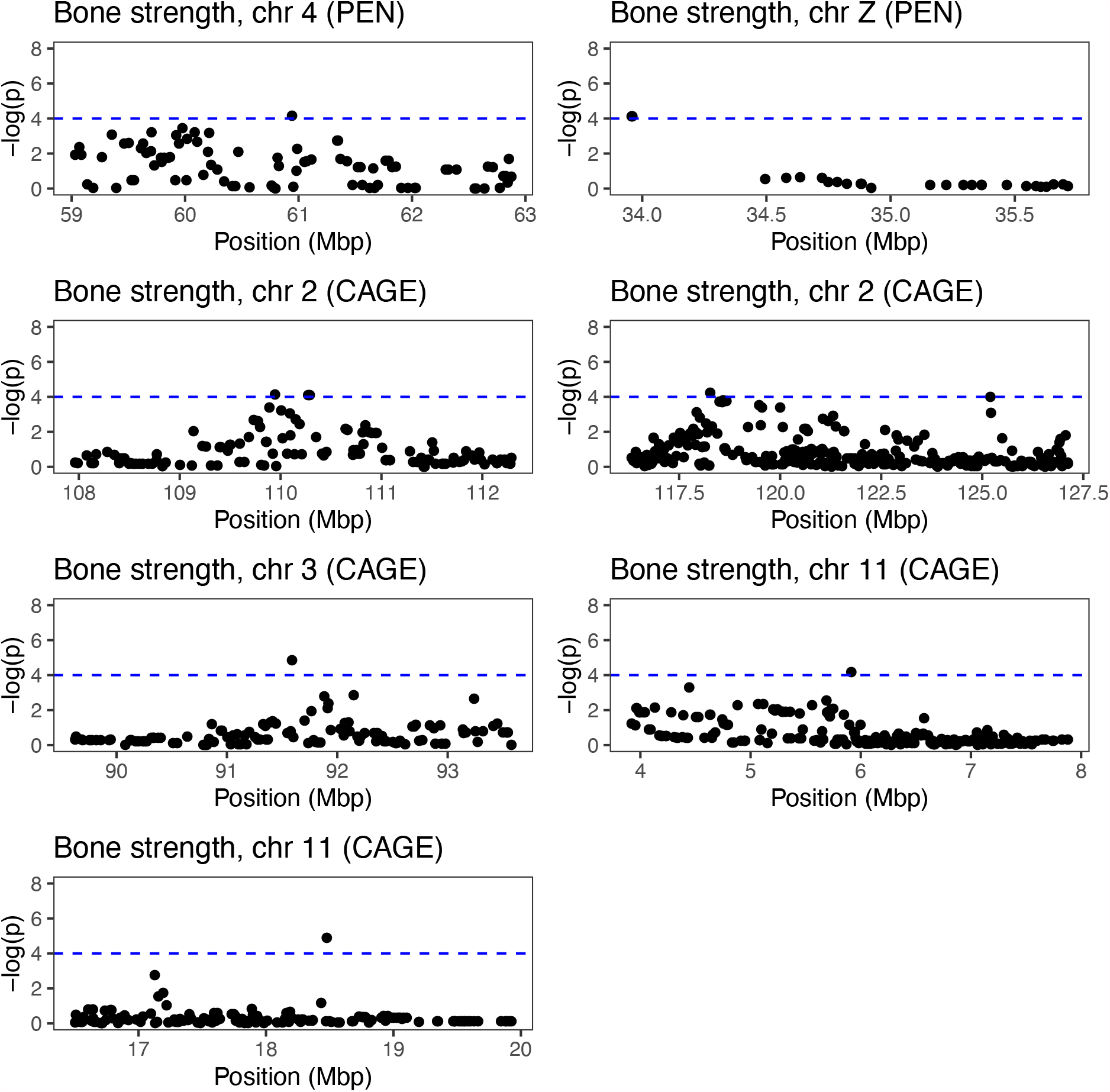
Zoomed-in view of suggestive genome-wide associations for bone breaking strength. The dashed blue line shows a suggestive threshold of 10^−4^.

**Supplementary Figure 8.**
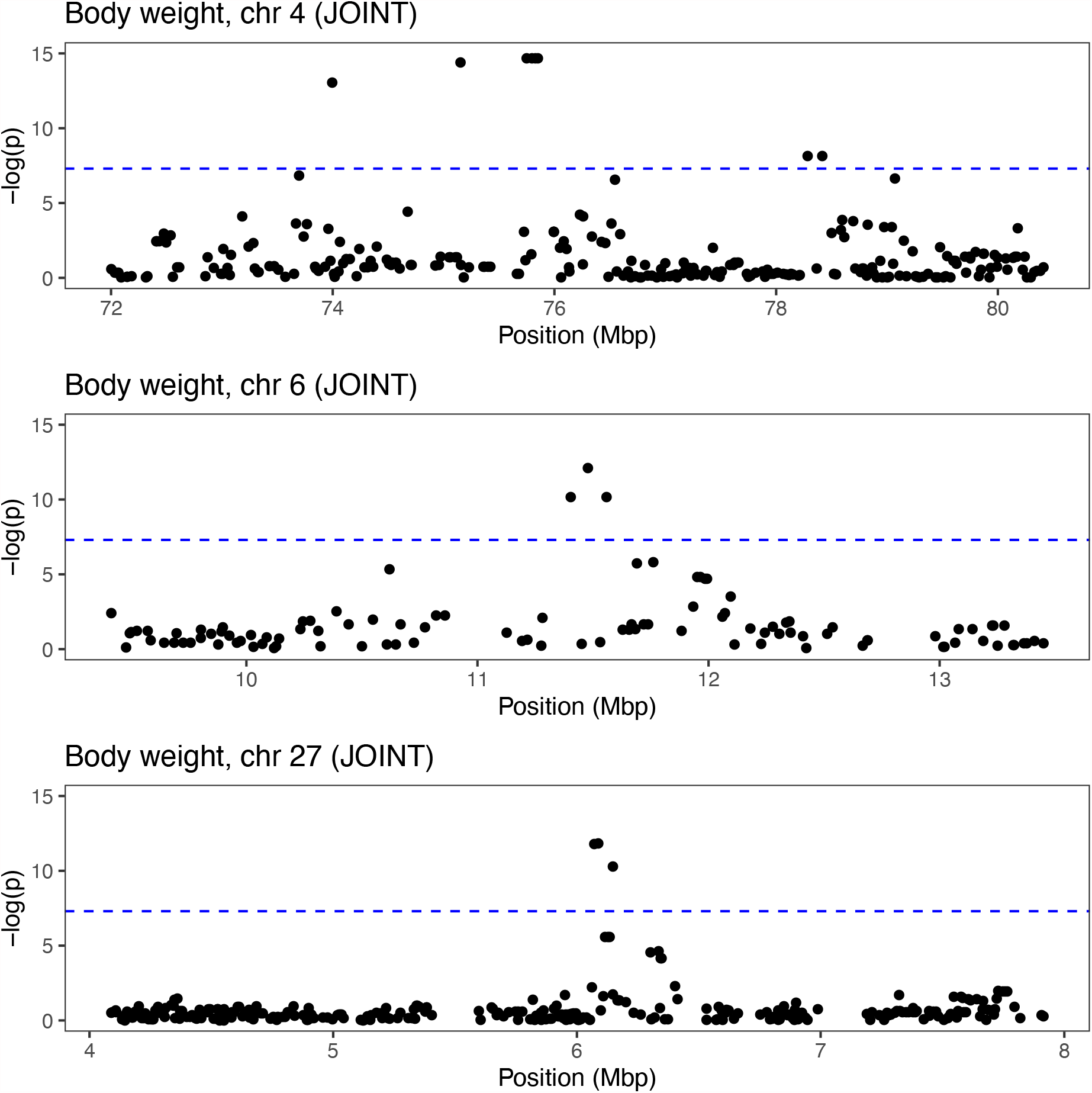
Zoomed-in view on genome-wide associations for body weight. The dashed red line shows a conventional genome-wide significance threshold of 5 * 10^−8^.

**Supplementary Figure 9.**
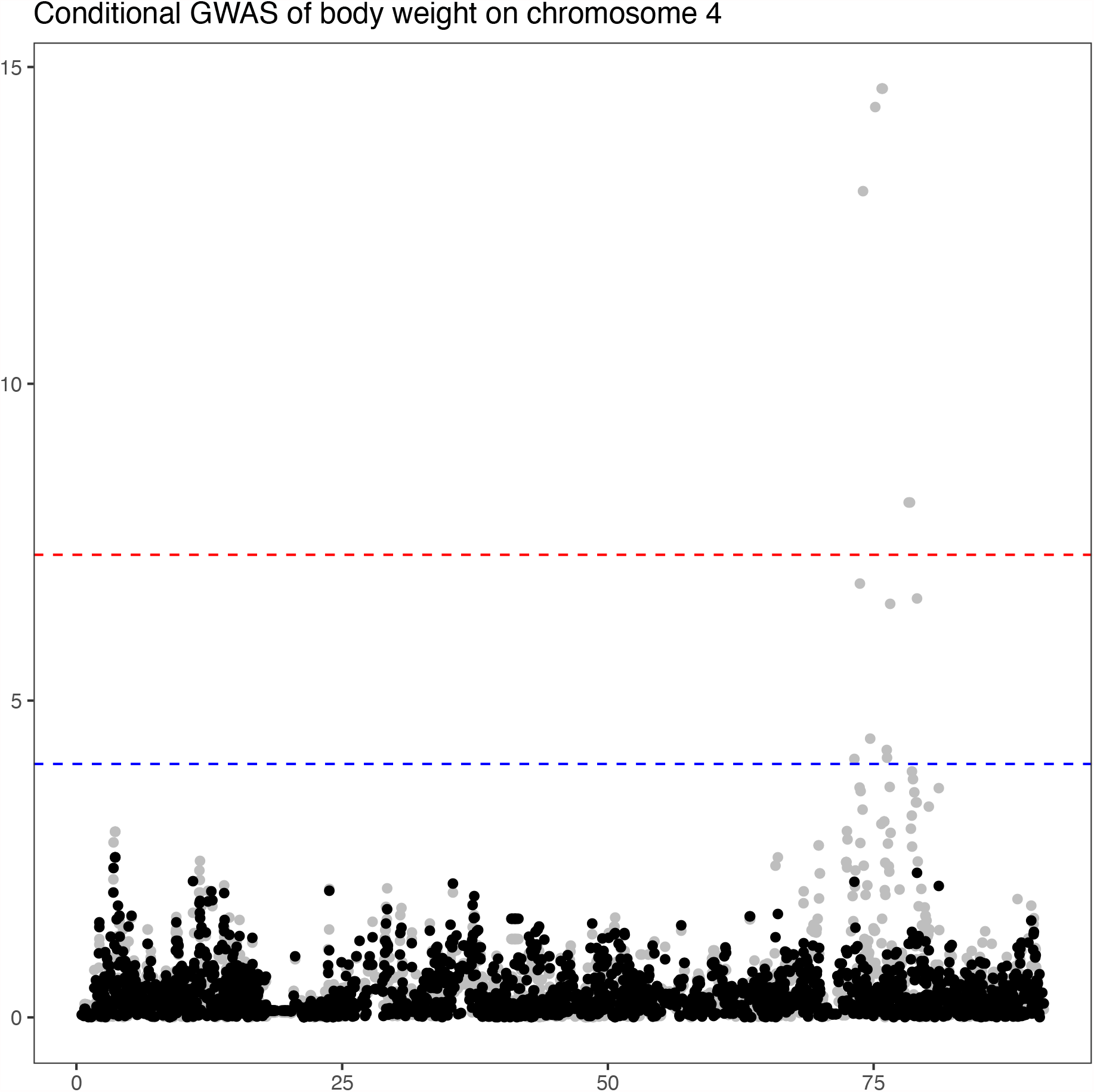
Conditional GWAS of the chromosome 4 locus for body weight. The plot shows the negative logarithm of the p-value for chromosome 4, with grey dots being the joint GWAS performed in the main analysis, and black dots a conditional GWAS including the lead SNP from the locus. This conditional scan removes associations throughout the region.

**Supplementary Figure 11.**
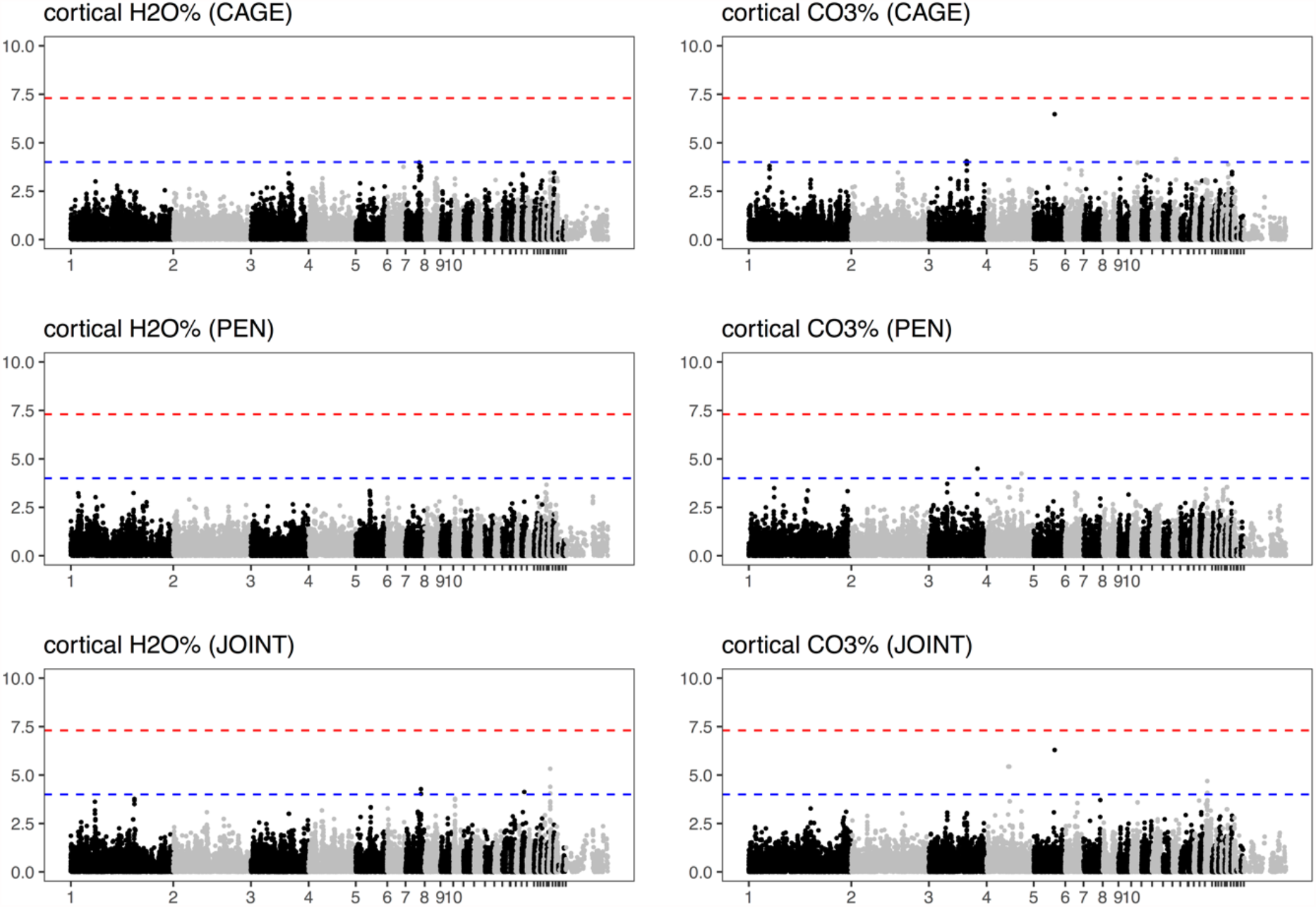
Genome-wide association of bone composition phenotypes that had significant heritability in both housing system. Chromosome names of the smaller chromosomes have been suppressed for legibility. The dashed red lie shows a conventional genome-wide significance threshold of 5 * 10^−8^, and the dashed blue line a suggestive threshold of 10^−4^.

**Supplementary Figure 12.**
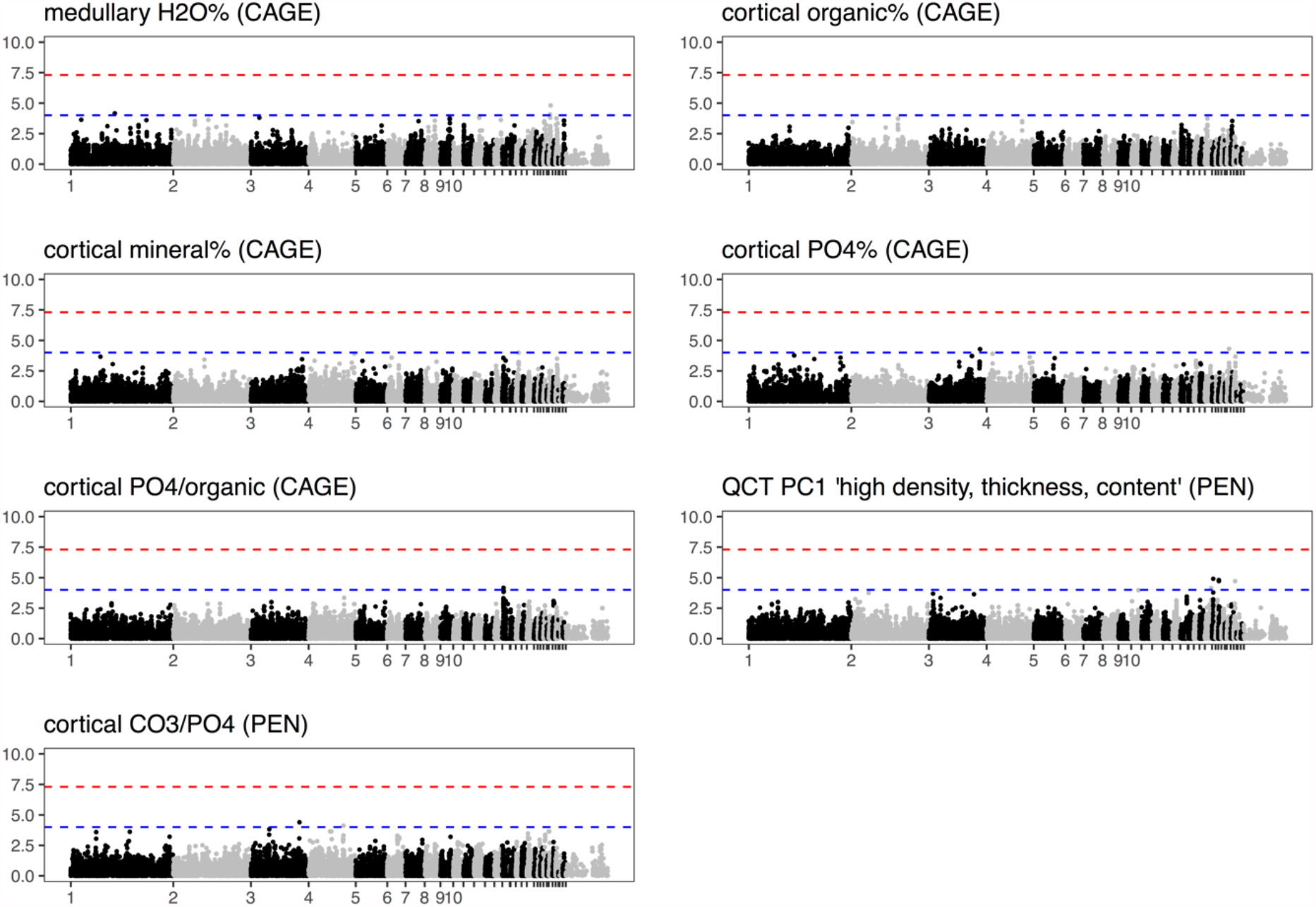
Genome-wide association of bone phenotypes that had significant heritability only in one housing system. Chromosome names of the smaller chromosomes have been suppressed for legibility. The dashed red line shows a conventional genome-wide significance threshold of 5 * 10^−8^, and the dashed blue line a suggestive threshold of 10^−4^.

**Supplementary Figure 13.**
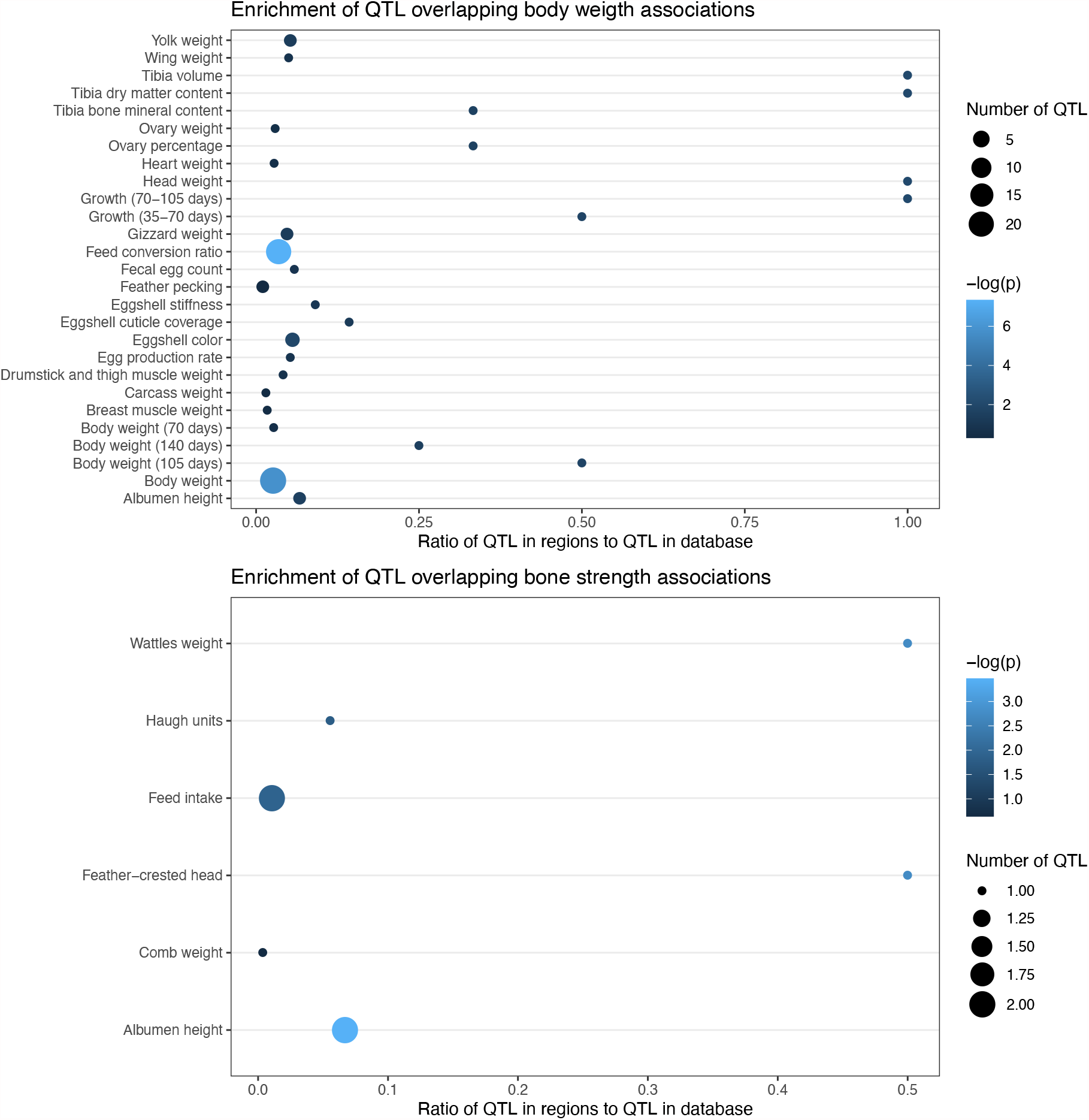
Enrichment of previously published QTL from the Chicken QTLdb database overlapping significant body weight and suggestive bone strength associations.

## Description of supplementary tables

Supplementary Table 1. Sample sizes.

Supplementary Table 2. Predefined candidate regions derived from previous studies.

Supplementary Table 3. Markers in predefined candidate regions with p < 0.01.

Supplementary Table 4. Genomic heritabilities and correlations from bivariate model.

Supplementary Table 5. Suggestive associations from genome-wide association studies.

Supplementary Dataset 1. Summary statistics for all markers from genome-wide association studies.

